# Intermittent Cytomegalovirus Infection Alters Neurobiological Metabolism and Induces Cognitive Deficits in Mice

**DOI:** 10.1101/2022.12.16.520745

**Authors:** Mark A.A. Harrison, Sara L. Morris, Grace A. Rudman, Daniel J. Rittenhouse, Chandler H. Monk, Siva SVP Sakamuri, MaryJane J. Jones, Md Mehedi Hasan, Mst Shamima Khatun, Hanyun Wang, Lucas P. Garfinkel, Elizabeth B. Norton, Chad Steele, Sangku Kim, Jay K. Kolls, S. Michal Jazwinski, Ricardo Mostany, Prasad VG Katakam, Elizabeth B. Engler-Chiurazzi, Kevin J. Zwezdaryk

**Author notes:** Correspondence (K.J.Z), (E.B.EC).

## Abstract

Risk factors contributing to dementia are multifactorial. Pathogens as risk factors for dementia is largely correlative with few causal relationships. Here, we demonstrate that intermittent cytomegalovirus (CMV) infection in mice, mimicking human chronic infection and reactivation/reinfection events, alters blood brain barrier (BBB) metabolic pathways. An increase in basal mitochondrial function is observed in brain microvasculature endothelial cells (BMEC) at 12 months post infection but not at earlier time points and is accompanied by elevated levels of superoxide, indicative of oxidative stress. Further, these mice score lower in cognitive assays as compared to age-matched controls. Our data show that repeated systemic infection with CMV, alters BBB metabolic function and impacts cognition. These observations provide mechanistic insights through which pathogens contribute to the progression of pathologies associated with dementia.

**In Brief:** Mechanistic evidence supporting an infectious etiology of dementia (e.g. Alzheimer’s Disease) are poorly defined. Harrison et al., show that intermittent infection with cytomegalovirus metabolically rewires the blood brain barrier and neighboring glial cells altering their function, resulting in decreased cognitive function.

## INTRODUCTION

Aging is a primary risk factor associated with cognitive decline and the development of dementia, such as Alzheimer’s disease (AD) (Hou et al., 2019; Moir et al., 2018). Therapeutic strategies have focused on extracellular amyloid-beta (Aβ) oligomers and soluble fragments, or intracellular hyperphosphorylated-tau tangles. However, these hallmarks of AD are not evident until the disease has been developing for years and, even at that point, patients are often in a pre-clinical phase of the disease with little to no cognitive symptoms (Dubois et al., 2016; Guzman-Velez et al., 2022). These findings suggest that other initiating or contributing factors exist which may be addressed decades prior to the development of any cognitive symptoms.

Pathogens are increasingly recognized as risk factors in the etiology of age-associated neurodegeneration (Harris and Harris, 2015; Moir et al., 2018). A single viral or bacterial infection in adult rodents has been shown to promote sickness behavior, induce systemic and central nervous system (CNS) inflammatory cascades, alter memory performance, and disrupted synaptic communication via neuronal loss (Hritcu et al., 2011; Kahn et al., 2012; Kranjac et al., 2012; McLinden et al., 2012; Tarr et al., 2011; Zhao et al., 2019). Humans do not experience a single infection during their lifetimes, but rather intermittent infections throughout life. The long-term consequences of intermittent infection are inadequately studied. Emerging preclinical evidence suggests that a higher lifetime infection burden impairs cognition, especially among transgenic mice carrying AD risk factors (Batista et al., 2019; Harris and Harris, 2015). Finally, studies in adult and aged humans show that higher cumulative exposure to HSV-1, cytomegalovirus (CMV), and bacterial infections was associated with impaired cognitive performance or an accelerated rate of cognitive decline (Dickerson et al., 2014; Katan et al., 2013; Lovheim et al., 2018; Lurain et al., 2013). Converging evidence indicates that pathogen exposure can promote features of dementia but the mechanisms for these effects have yet to be elucidated.

In this study, CMV was used a as a model pathogen. CMV is a ubiquitous herpesvirus exhibiting 40-80% seropositivity in the human population (Hoehl et al., 2020). Common to all herpesviruses, following acute infection, CMV becomes latent and reactivates sporadically throughout the lifetime of the host resulting in intermittent infection. Further, different CMV strains are thought to infect individuals throughout life. CMV’s ability to evade immune clearance enables a persistent phenotype that is associated with immunosenescence, inflammaging, and memory T cell inflation (Holtappels et al., 2000). Most individuals seroconvert before the age of 40 suggesting that the cycle of latency/reactivation or reinfection continues for decades (Bate et al., 2010).

The role of CMV in unhealthy aging, including dementias, remains largely correlative. Data suggesting individuals positive for CMV have an increased risk for AD and accelerated cognitive decline (Barnes et al., 2015; Lee et al., 2020) are contrasted with data suggesting a weak association (Warren-Gash et al., 2019). Ultimately, a lack of mechanisms linking pathogens and the origins of dementia impede progress. Intriguingly, CMV pathogenesis mirrors many of the features of AD pathology including calcium dysregulation (Sharon-Friling et al., 2006), amyloid deposition (Lurain et al., 2013), iron dysfunction (Crowe et al., 2004), autophagic (Taisne et al., 2019), and mitochondrial dysfunction (Betsinger et al., 2021; Combs et al., 2020; Harrison et al., 2022).

The ability of CMV to alter the metabolic profile of host cells to produce the biomaterials and energy needed for viral replication is well described (Combs et al., 2020; Harrison et al., 2022; Saffran et al., 2007; Spaggiari et al., 2008; Suliman et al., 2004; Vastag et al., 2011). This metabolic reprogramming often results in impaired bioenergetics and subsequent increases in reactive oxygen species (ROS) production. Interestingly, ROS production and impaired metabolism have been implicated in numerous neurodegenerative diseases (Cai et al., 2012; Kim et al., 2015; Li et al., 2013; Muddapu et al., 2020). Additionally, the Mitochondrial Cascade Hypothesis posits that AD pathogenesis is driven by progressive mitochondrial dysfunction (Ashleigh et al., 2022). Further, it has been shown that mitochondrial function directly alters the secretion or storage of Aβ (Wilkins et al., 2022). Finally, studies have demonstrated that metabolic and mitochondrial changes alter cellular proteostasis and precede the onset of clinical symptoms of AD (Swerdlow, 2020). Therefore, CMV, with its potent capacity to reprogram cellular metabolism is an ideal candidate for assessing the impact of pathogens on aging-associated pathology such as AD.

In this study, we demonstrate that intermittent CMV infection induces metabolic alterations in the blood brain barrier (BBB) that correlate with decreases in cognitive function. We observe that endothelial cells isolated from the BBB display enhanced oxidative phosphorylation (OXPHOS) and consequently produce elevated levels of superoxide (SO) following the pro-inflammatory immune response. These findings suggest that infection frequency and burden may accelerate cognitive decline by modulating BBB metabolism and increasing oxidative stress.

## RESULTS

### Intermittent MCMV infection exhibits delayed inflammatory cytokine expression in the brain

Pathogen exposure and burden increase with age (Kline and Bowdish, 2016). Using a reductionist model, we examined how a single herpesvirus (CMV) can contribute to accelerated hallmarks of aging and dementia. As CMV exhibits species tropism, we used the well described Smith strain of murine CMV (MCMV) in our studies(Fisher and Lloyd, 2020). We employed a murine model that integrates intermittent infection and natural aging starting with young adult mice aged 8 weeks (Figure 1A). In immunocompetent individuals, human CMV infection is asymptomatic, or results in mild flu-like symptoms and general malaise. Our murine model behaves similarly, exhibiting slight but significant changes in body weight 2-3 days post infection (DPI), but rapid recovery equaling mock-infected mice by 5 DPI (Figure 1B). Repeat MCMV exposures at 3-, 6-, and 9-months post infection (MPI) do not result in significant changes in body weight (Figure S1A). Additionally, there were no significant changes in weight at 1 MPI (Figure 1C) and mice exposed to intermittent infection did not exhibit significant changes in weight loss over 12 months (Figure S1B). Plasma IgM specific to MCMV exhibited high concentrations 7 DPI after initial infection (Figure 1D) but were reduced by 1 MPI. Subsequently, MCMV specific IgM concentrations increased with reinfection. MCMV specific IgG concentrations correlated strongly with MCMV specific IgM plasma concentration and significantly increased as a function of challenge number and remained quantitively similar from 6-12 MPI (Figure 1E). All mock-infected mice failed to elicit an antibody response against MCMV, confirming the infection status remained negative throughout our experimental timeline. To validate our measured results were not the result of active viral shedding, PCR analysis was completed using salivary glands. MCMV can be detected at high viral loads in the salivary gland (Campbell et al., 2008). At 7 DPI we observed high levels of UL123, a viral gene expressed early during replication. Expression of UL123 decreases by 1 MPI (Figure S1C). There was no replicating MCMV detected in salivary glands at 6- or 12-month timepoints suggesting a functional immune response and establishment of MCMV latency.

**Figure 1:**
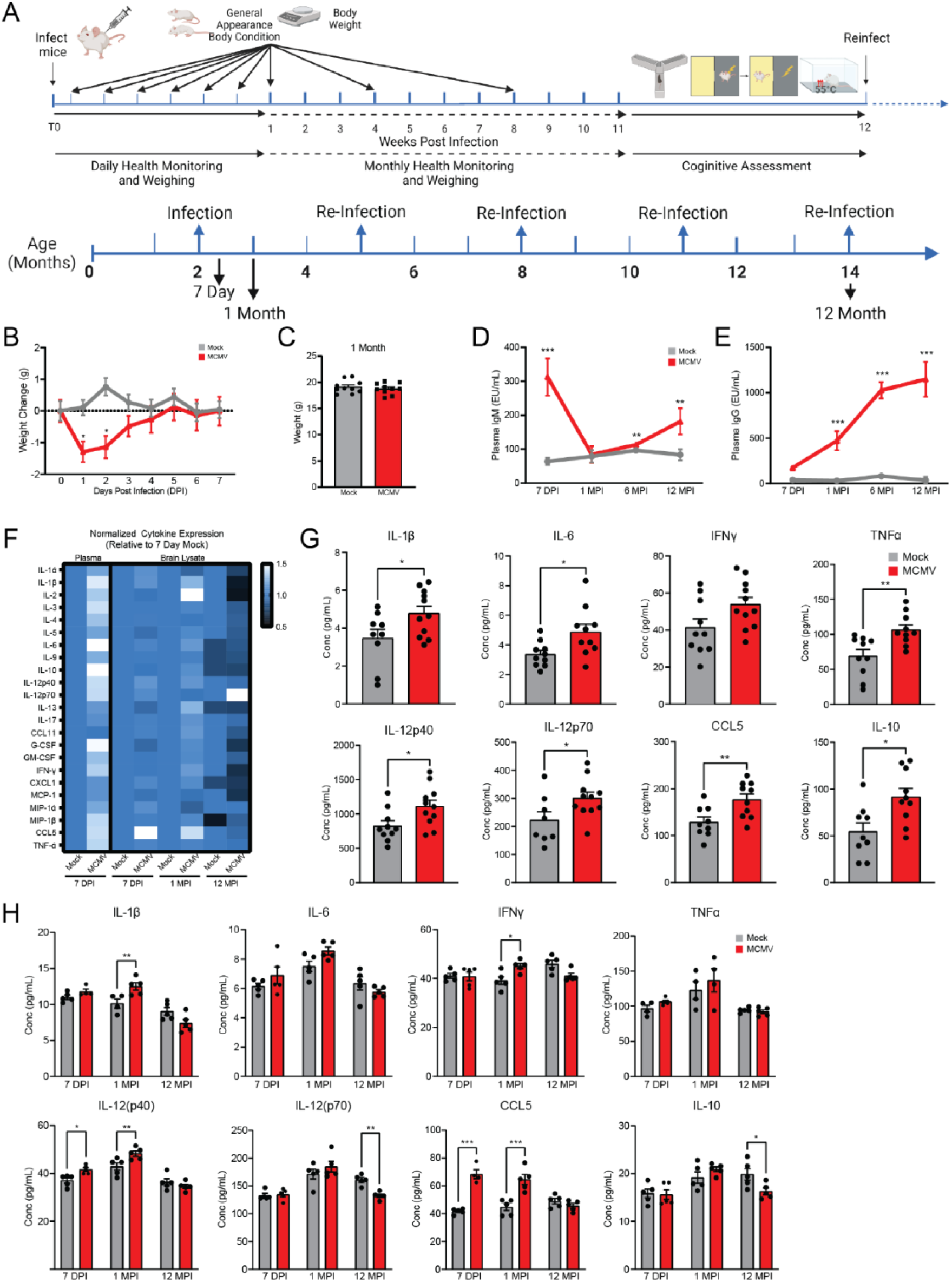
Intermittent MCMV infection exhibits delayed inflammatory cytokine expression in the brain. (A) Schematic outlining the experimental design and re-infection timeline. (B) Weight change in mice over 1 week following initial infection. (C) Weight of mock-(grey) and MCMV-infected (red) mice at 1 month post infection. ELISA-derived concentration of MCMV-specific (D) IgM and (E) IgG from plasma across time. (F) Heatmap of cytokine expression in plasma and brain homogenate at 7 DPI, 1 MPI, and 12 MPI normalized to 7 DPI mock infected measurements. Levels of IL-1β, IL-6, interferon IFNγ, TNFα, IL-12p40, IL12-p70, CCL5, and IL-10 from (G) plasma at 7 DPI (H) and brain homogenate at 7 DPI, 1 MPI, and 12 MPI as measured by a multiplex immunoassay. Graphs represent pooled data from at least three individual animals. Mean ± the SEM. *, *p* < 0.05; **, *p* < 0.01; ***, *p* < 0.001. DPI = days post infection, MPI = months post infection. EU = ELISA units.

Previous reports have suggested that in MCMV models, pro-inflammatory cytokine responses may inhibit the innate immune response contributing to viral pathogenesis (Stacey et al., 2017; Stacey et al., 2011). We observe differential expression of cytokines in relation to infection frequency, time, and the origin of the sample (Figure 1F). In blood at 7 DPI, we observed significant upregulation of pro-and anti-inflammatory cytokines including IL-1β, IL-6, TNFα, IL-12p40, IL-12p70, CCL5, and IL-10 as compared to mock infected animals. (Figure 1G, Figure S1D). CCL5 has previously been identified as a commonly increased chemokine following CMV infection (Caldeira-Dantas et al., 2018; Wang et al., 2004). In contrast, at 7 DPI, we observed mild pro-inflammatory cytokine increases in brain homogenate that was limited to IL-12(p40) and CCL-5 (Figure 1H, Figure S1E). Interestingly, at 1 MPI, we observed an elevated expression of inflammatory cytokines similar to that measured in the periphery at 7 DPI (Figure 1H, Figure S1E). At this timepoint, the inflammation in the periphery was resolved (Figure S1F). However, at 12 MPI, after 4 MCMV challenges, inflammatory cytokine levels in the brain are consistently lower in expression (Figure 1H, Figure S1E) compared to the mock infected controls. This pattern of cytokine expression was not observed in mock-infected mice (Figure S1G). Based on these findings, we conclude that peripheral inflammation results in a delayed inflammatory response in the CNS.

### Intermittent MCMV infection alters the transcriptional profile of brain cells

We next used Multiome analysis (simultaneous single cell gene expression and ATAC-seq) to examine transcriptional changes in various cell populations. Whole brain lysate from mock-infected 12 MPI animals was compared to 12 MPI animals that had experienced intermittent CMV infection. Dimensionality reduction analysis using UMAP showed clustering of 7 distinct cell populations based on single-cell RNA expression (Figure 2A), single-cell chromatin accessibility (Figure 2B) and integrated scATAC/scRNA (Figure 2C). To confirm observations from single cell transcriptional data, we assessed brain cells phenotypically and metabolically using spectral flow cytometry (Table S1, Figure S2A). At all timepoints examined, there were minor changes in the percent of brain cell populations (Figure 2E, 2F, 2G). We observed significant increases in T-cell populations in infected mice at all time points, indicating enhanced extravasation from the bloodstream into the neural parenchyma. Of the other cell types assayed, there were no significant changes between Mock and MCMV infected animals at any timepoint. However in young adult mice, at 7 DPI, we observed increased proliferation rates in activated microglia/macrophages and T cells (Figure 2H). These were accompanied by decreased proliferation of brain microvascular endothelial cells (BMECs) (Figure 2H). Taken together, these findings demonstrate expanding immune populations in response to initial infection or the subsequent inflammatory response. T-cell populations exhibited changes in proliferation rate, as measured by Ki-67 positivity, with a decrease at 1 MPI and an increase at 12 MPI respectively (Figure 2I, 2J).

**Figure 2:**
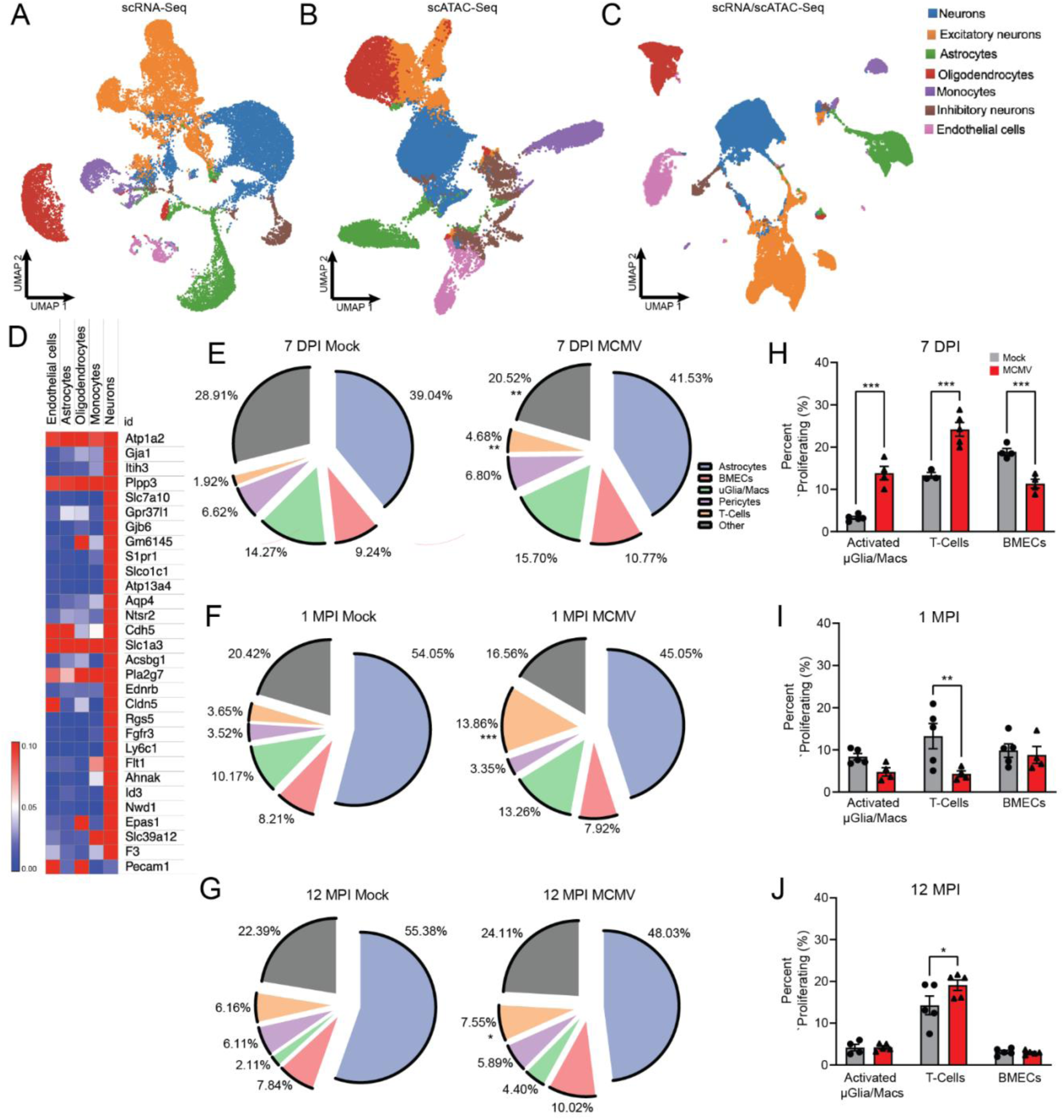
Intermittent MCMV infection alters the transcriptional profile of brain cells. Multiome libraries were generated and integrated. UMAP clustering of cell types based on single cell expression (A), chromatin accessibility (B) and integrated scATAC/scRNA (C). (D) Heatmap of transcriptional changes plotted by cell type. Population distributions as a percentage of live cells isolated in mock- and MCMV-infected mice at (E) 7 DPI, (F) 1 MPI, (G) 12 MPI as assessed by flow cytometry. Percentage of proliferating cells in different cell populations as measured by Ki-67 positivity at (H) 7DPI, (I) 1 MPI, and (J) 12 MPI. Graphs represent pooled data from at least three individual animals. Mean ± the SEM. *, *p* < 0.05; **, *p* < 0.01; ***, *p* < 0.001.

### Blood brain barrier metabolic pathways are altered with increased MCMV exposure

Our current data does not suggest that MCMV in our model directly infects the brain. Others have reported similar findings (Reuter et al., 2004). If peripheral MCMV infection can influence changes in the brain, the initial site of interaction is likely the BBB. Our Multiome data showed significant changes in transcriptional expression of brain endothelial cells associated with metabolism (Slc1a3 and Atp1a2) and BBB function (Cdh5, Cldn5, plpp3, Pla2g7, and Pecm1) (Figure 2D). Due to these findings, we focused on characterizing the metabolic profile of BMECs. We used glucose transporter 1 (GLUT1), hexokinase 2 (HKII), translocase of the outer mitochondrial membrane 20 (TOMM20) and voltage dependent anion channel 1 (VDAC1) as proxies for glycolytic and OXPHOS pathways (Figure 3A). In our characterization of BMECs we identified a large population that had high expression of eNOS while a minority exhibited low eNOS expression. Interestingly, in MCMV infected animals at 7 DPI, both total and eNOS low BMECs had elevated expression of GLUT1 relative to mock infected animals (Figure 3B). This is contrasted with a downregulation of GLUT1 in total BMECs (Figure 3C) and downregulation in the eNOS high group at late timepoints, as compared to the mock infected controls (Figure 3D). Hexokinase expression at 7 DPI was unchanged (Figure S3A), decreased at 1 MPI (Figure S3B), and only increased at 12 MPI in the eNOS low group (Figure S3C). Taken together, initial infection appears to enhance glycolytic function, but that change is transient and trends towards decreased glycolysis as a function of repeat infection and age. In contrast, mitochondrial pathways trended towards enhancement in young adult mice at 7 DPI as indicated by VDAC1 (Figure 3E) and TOMM20 (Figure S3D) expression. At 1 MPI VDAC1 expression (Figure 3F) is decreased and TOMM20 (Figure S3E) remains unchanged. By 12 MPI, total BMECs and the eNOS high group have significantly upregulated VDAC1 expression which suggests increased OXPHOS function as VDAC1 is the primary channel for ATP/ADP exchange in mitochondria (Figure 3G). Expression of the protein transporter TOMM20 was upregulated in eNOS high groups at 7 DPI (Figure S3D), unchanged at 1 MPI (Figure S3E), and downregulated at 12 MPI (Figure S3F). Previous research has demonstrated that, despite having access to sufficient oxygen, BMECs are highly glycolytic (McDonald et al., 2022). The young adult mice, at 7 DPI, protein expression results suggest that MCMV enhances this metabolic preference. However in aged mice, by 12 MPI, these cells have been reprogrammed to enhance OXPHOS as indicated by VDAC1 protein expression. The impact of these changes is unknown but they suggest altered BMEC metabolic function and possibly BBB integrity.

**Figure 3:**
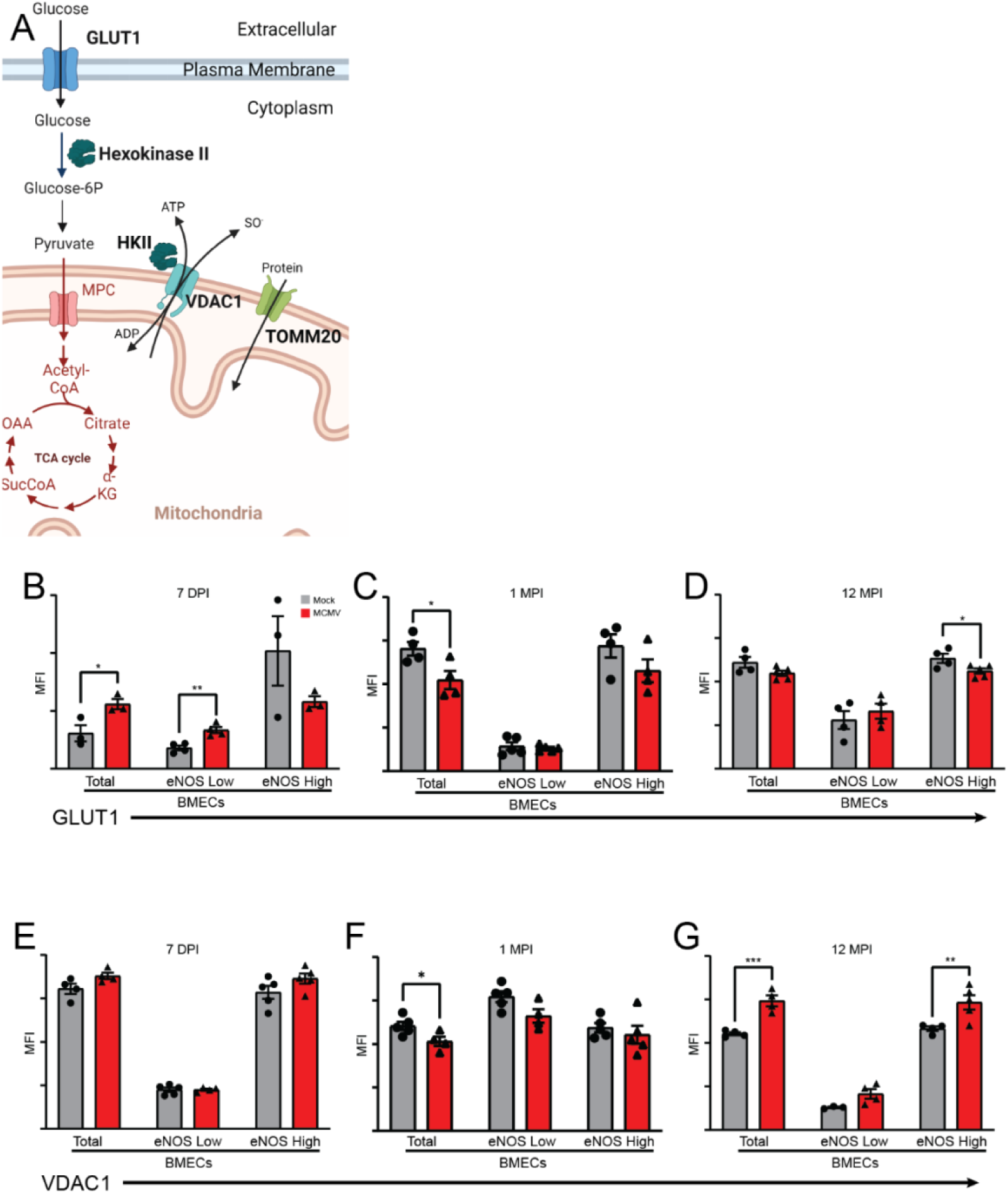
Blood brain barrier metabolic pathways are rewired with increased MCMV exposure. (A) Cartoon identifying key proteins in glycolysis and mitochondrial function. Flow cytometry was used to examine expression of GLUT1 in populations of BMECs at (B) 7 DPI, (C) 1 MPI, and (D) 12 MPI. Expression of VDAC1 in populations of BMECs at (E) 7 DPI, (F) 1 MPI, and (G) 12 MPI. Graphs represent pooled data from at least three individual animals. Mean ± the SEM. *, *p* < 0.05; **, *p* < 0.01; ***, *p* < 0.001. eNOS = endothelial nitric oxide synthase, MFI = mean fluorescent intensity.

### Hallmarks of mitochondrial dysfunction and oxidative stress are elevated in BMECs intermittently exposed to MCMV

To better understand the impact of altered expression of metabolic markers, we isolated BMECs from the BBB and completed Seahorse bioanalyzer analysis (Figure 4A). At 1 MPI there were no significant changes to OXPHOS including mitochondrial basal respiration, proton leak, non-mitochondrial respiration or mitochondrial ATP production rate (Figure 4B, 4C). Similarly, there were no changes in maximal respiration, spare respiratory capacity or total ATP production rate between mock and infected mice (Figure S4A). Intriguingly in the aged mice, 12 MPI, the basal respiration rate, the minimal work required by the mitochondria to meet metabolic demand, was significantly increased in MCMV-infected mice (Figure 4D, 4E). This suggests that BMEC mitochondria from infected mice work harder to attain the same level of metabolic homeostasis as BMECs from mock-infected mice. No changes in maximal respiration or spare capacity (Figure S4B) were observed. Critically, we measured significant increases in proton leak and non-mitochondrial respiration (Figure 4E) in BMEC from MCMV infected mice, suggesting the presence of oxidative and mitochondrial stress. Mitochondrial ATP production rate was not found to be increased (Figure 4E). To validate our findings, live cell examination of mitochondria was performed by flow cytometry on CD31^+^ cells isolated from the brain. No changes in mitochondrial mass were detected at 1 MPI (Figure 4F) but increases were observed in aged mice, 12 MPI (Figure 4G), when comparing mock- vs MCMV-infected. Critically, mitochondria-specific superoxide was significantly upregulated only in mice exposed to intermittent infection (Figure 4H, Figure 4I). Together, these data suggest intermittent MCMV infection decreases BMEC mitochondrial efficiency and increases oxidative stress.

**Figure 4:**
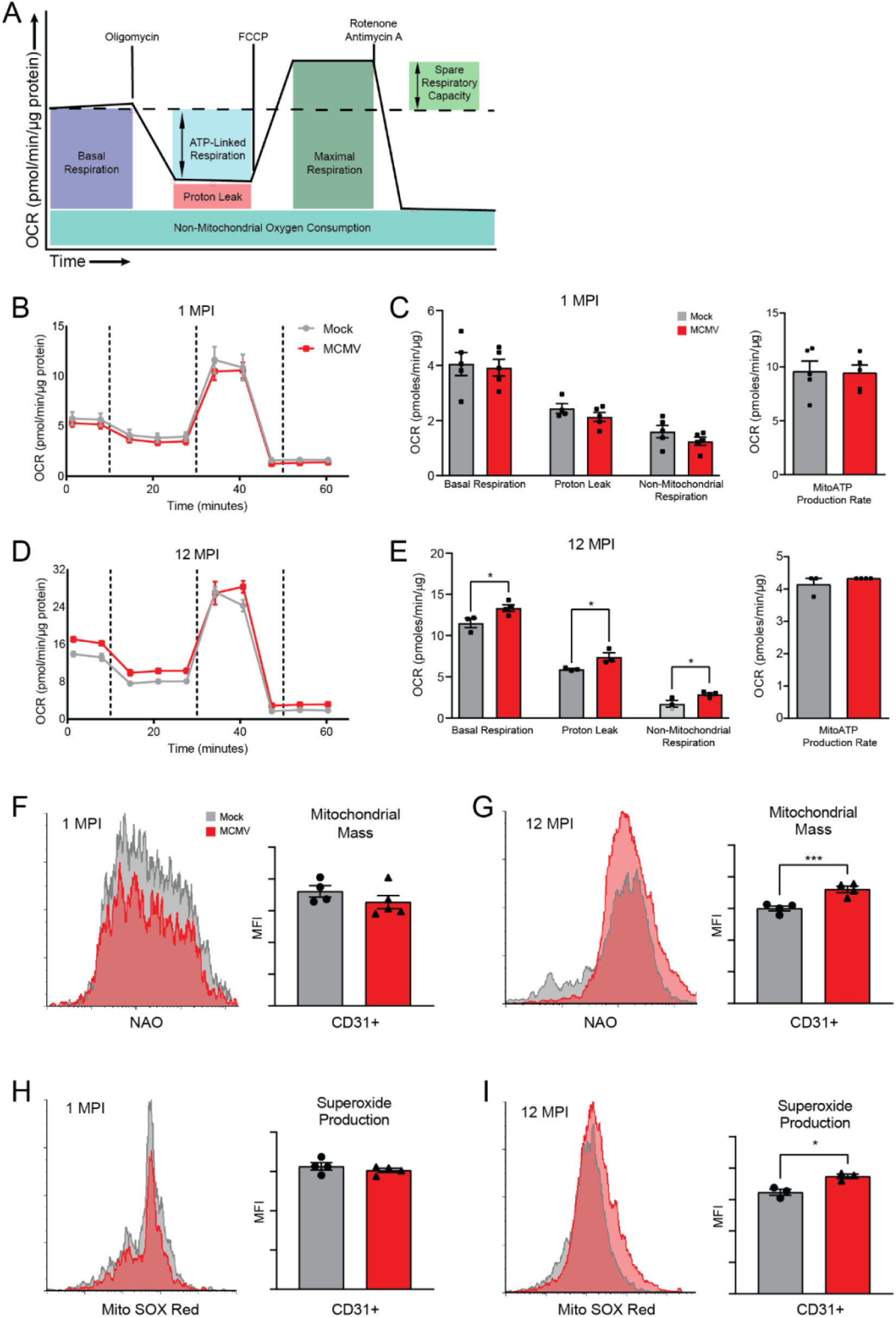
Hallmarks of mitochondrial dysfunction and oxidative stress are elevated in BMECs intermittently exposed to MCMV. (A) Cartoon illustrating how mitochondrial function was measured in BMECs using the Seahorse XFe24 MitoStress Kit. All samples were normalized to protein concentration. Representative data output of BMECs from the (B) 1 MPI and (D) 12 MPI timepoints. Basal Respiration, Proton Leak, Non-Mitochondrial Oxygen Consumption and mitochondrial ATP production rate were derived from the (C) 1 MPI and (E) 12 MPI measurements. Mitochondrial mass was determined by flow cytometry at (F) 1 MPI and (G) 12 MPI using nonyl acridine orange (NAO). Superoxide production was assessed using mitoSOX Red at (H) 1 MPI and (I) 12 MPI. Graphs represent pooled data from at least three individual animals. Mean ± the SEM. *, *p* < 0.05; **, *p* < 0.01; ***, *p* < 0.001. OCR = oxygen consumption rate, MPI = months post infection.

### MCMV infection metabolically rewires astrocytes as a function of age

We next asked if metabolic changes were restricted to the BBB or if additional changes were occurring in the brain parenchyma. Due to their close interactions and functional cooperation with the BBB, astrocytes were further examined. In addition to total astrocytes we also further subdivided this group into their pro-inflammatory (A1) and anti-inflammatory (A2) phenotypes. Astrocytes uptake metabolites and proteins from the BBB to supply nutrients to neurons (Figure 5A). Global decreases in astrocytic metabolic function, both glycolysis and OXPHOS, have been associated with AD and result in insufficient metabolic support for neighboring neurons (Andersen et al., 2022). GLUT1 in astrocytes is responsible for uptake of glucose from the bloodstream (Zlokovic, 2011) and AD patients have lower glucose levels in the brain (Herholz et al., 2002; Marcus et al., 2014; Mosconi et al., 2010). In young adult mice, 7 DPI and 1 MPI, GLUT1 expression in astrocytes is unchanged between mock and MCMV-infected mice but decreases in aged mice, 12 MPI (Figure 5B). HKII levels are significantly decreased at 1 MPI but are unchanged at 12 MPI (Figure S5A).

**Figure 5:**
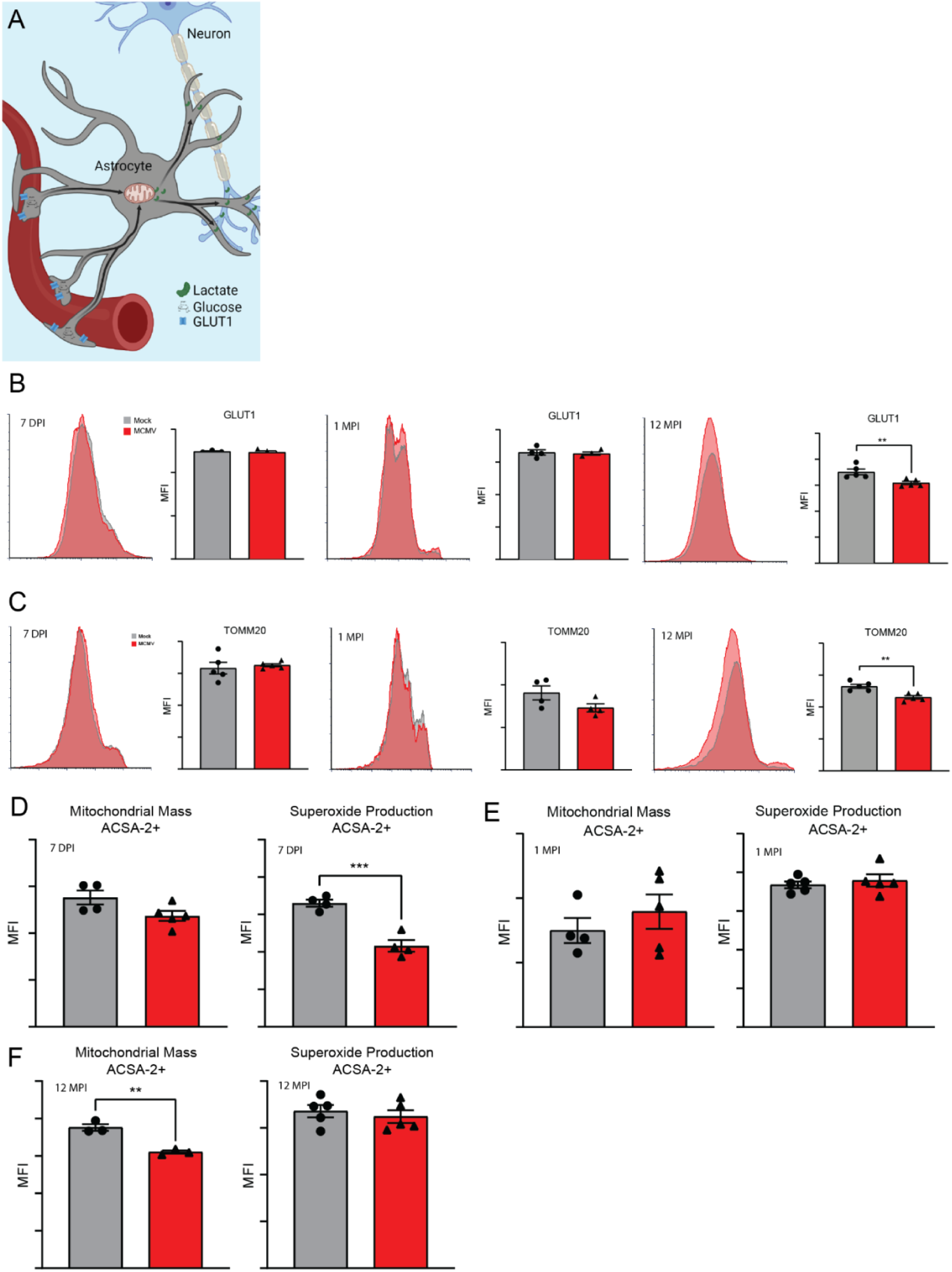
MCMV infection metabolically rewires astrocytes as a function of age. (A) Cartoon demonstrating the flow of metabolites from the bloodstream through astrocytes to neurons. Flow cytometry expression of (B) GLUT1 and (C) TOMM20 in astrocytes at 7 DPI, 1 MPI and 12 MPI. Mitochondrial mass and superoxide production were determined using nonyl acridine orange and mitoSOX red staining respectively and are quantified at (5D) 7 DPI, (5E) 1 MPI, and (5F) 12 MPI. Graphs represent pooled data from at least three individual animals. Mean ± the SEM. *, *p* < 0.05; **, *p* < 0.01; ***, *p* < 0.001. DPI = days post infection, MPI = months post infection. ACSA-2 = astrocyte cell surface antigen 2.

OXPHOS function, as indicated by the protein import channel subunit TOMM20, demonstrated a significant decrease in expression at 12 MPI in the MCMV-infected mice. TOMM20 expression is unchanged in A1 astrocytes at 7 DPI but increases in A2 astrocytes (Figure S5B). However, in aged mice, 12 MPI, total astrocytes showed decreased TOMM20 expression (Figure 5C), findings that repeated in A1 and A2 populations (Figure S5C). These findings suggest that mitochondrial function is altered due to infection burden. No changes in VDAC1 were observed (Figure S5D). Mitochondrial mass did not change at 7 DPI in ACSA-2 positive cells but there was significantly less superoxide production in MCMV-infected animals (Figure 5D). At 1 MPI both mass and superoxide production were similar in both groups of animals (Figure 5E). By 12 MPI the mitochondrial mass of astrocytes in the MCMV-infected mice was significantly decreased, however they were producing similar levels of superoxide (Figure 5F). These findings agree with the decreases in TOMM20; however the similar levels of superoxide suggest increased oxidative stress. Our results suggest that different cell populations in the brain, exhibit distinct metabolic changes in response to intermittent CMV infection and age.

### Increased T cell accumulation in the brain

An increased percent of live T cells was detected in both young adult, 7 DPI and 1 MPI, and aged mice, 12 MPI, indicating increased extravasation from the vasculature (Figure 2E, 2F, 2G, 6A, and 6B). In the 1 MPI cohort, T cells had decreased expression of OXPHOS and glycolysis metabolic markers relative to mock infected animals (Figure 6A). In the 12 MPI cohort, T cells had higher expression of OXPHOS and glycolysis metabolic markers relative to mock infected animals (Figure 6B). Additionally, these cells had significantly decreased H3K27me3 expression suggesting a more transcriptionally active population. The highly bioenergetic nature of these cells and retention in the parenchyma despite the absence of active infection or inflammation suggest that they represent a population of tissue resident T memory cells. Previous literature using a similar MCMV model described brain resident memory CD8 T cells acting as inhibitors of MCMV reactivation (Brizic et al., 2018). The presence of resident memory T cells in the CNS may contribute to disease progression as the CNS has conventionally been perceived as an immune privileged environment.

**Figure 6:**
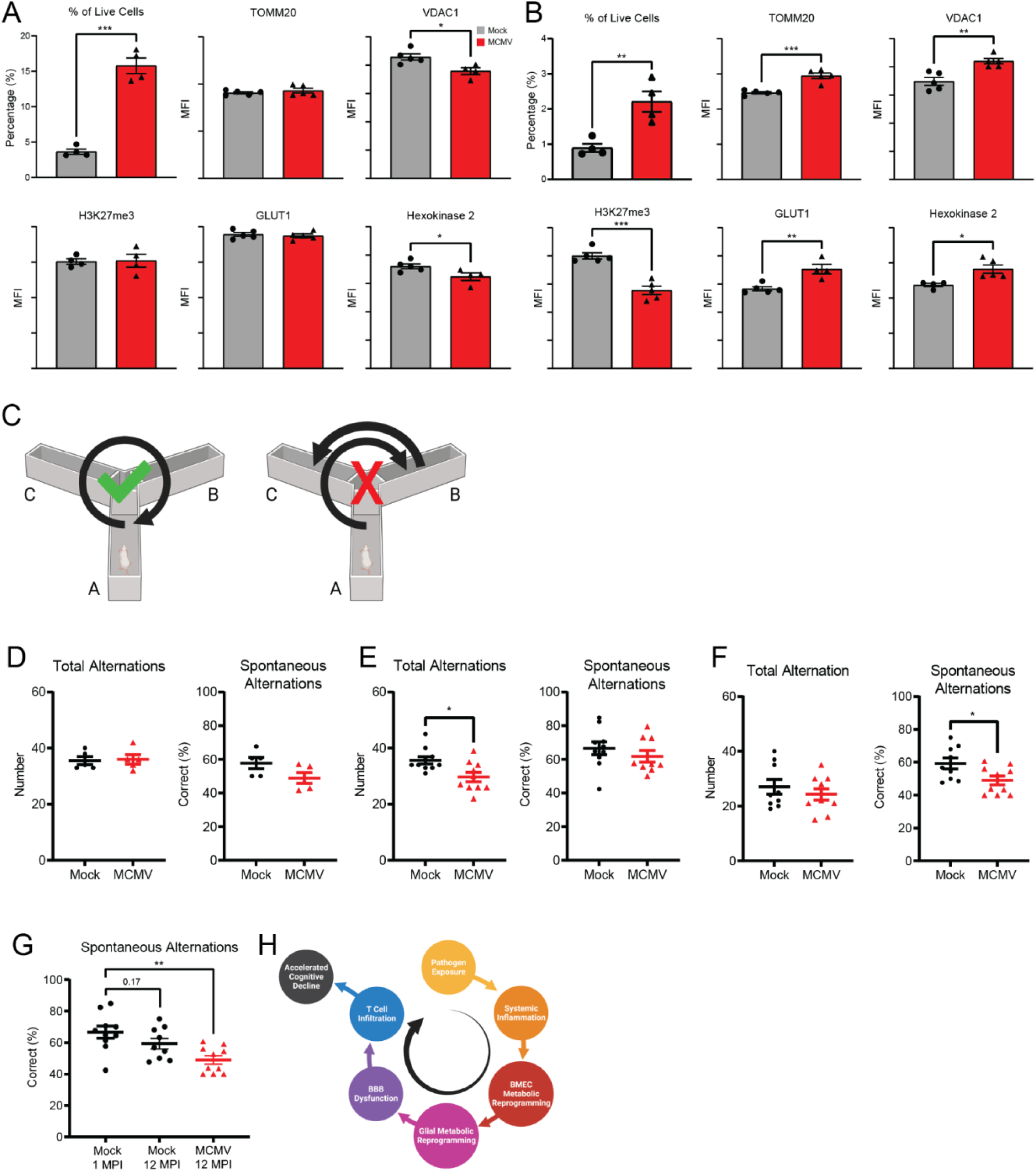
Cognition is impaired with intermittent MCMV infection. Proliferation, epigenetic, and metabolic markers of CNS resident T cells were examined using flow cytometry at (A) 1 MPI and (B) 12 MPI. (C) Schematic describing correct versus incorrect alternations in the Y-maze. Quantification of number of alternations and percentage of spontaneous correct alternations at (D) 7 DPI, (E) 1 MPI, and (F) 12 MPI mice. (G) Comparison of spontaneous alternations examining the interactions between age and infection status. (H) Diagram presenting the model of pathogen associated accelerated cognitive aging. Graphs represent pooled data from at least three individual animals. Mean ± the SEM. *, *p* < 0.05; **, *p* < 0.01; ***, *p* < 0.001. MPI = months post infection, MFI = mean fluorescent intensity.

### Cognition is impaired with intermittent MCMV infection

To understand the physical manifestations of our collective molecular results, we completed cognitive assays on all mice used in the experiments described in this manuscript. Hot plate (Figure S6A) and passive avoidance (Figure S6D) assays were completed to identify innate differences in nociceptive ability and fear-aggravated learning. At 1- (Figure S6B) and 12 MPI (Figure S6C), both groups exhibit similar latency to behavior and number of behaviors demonstrating that their nociceptive capabilities are intact and similar between groups (Woolfe and Macdonald, 1944). At 1- (Figure S6E) and 12 MPI (Figure S6F), both groups exhibit near perfect retention of learning after 24 hours. This is indicative that fear-based learning events (shock) with high salience are still able to be learned regardless of infection status. In young adult mice, 7 DPI, Y-maze spontaneous alternation assessment of short-term recognition memory (Figure 6C) demonstrated very similar number of alternations and spontaneous, correct alternations (Figure 6D). At 1 MPI, MCMV-infected mice exhibited a significantly lower number of total alternations, but no significant changes to spontaneous alternations (Figure 6E). Interestingly, in aged mice, 12 MPI (3 months after last infection), no difference in total alternations was measured but there was a significant decrease in spontaneous alternations, indicative of impaired spatial working memory (Figure 6F).

Additionally, we observed a trend towards decreased spontaneous alternations when comparing young adult, 1 MPI, and aged, 12 MPI, mock-infected mice (Figure 6G). These findings are in agreement with previous studies of cognitive impairment in BALB/cJ mice in which cognitive decline was first measurable at 12 months of age (Esquivel et al., 2020). We see a significant decrease in cognition with infection status and age when comparing 1 MPI mock and 12 MPI MCMV-infected cohorts. Collectively, our results suggest that increased viral burden, metabolically reprograms BMEC and glial metabolism, driving BBB dysfunction, T cell infiltration and cognitive decline (Figure 6H). This suggests synergy between viral burden and age as they relate to cognitive decline.

## DISCUSSION

Dementia, including AD, are identified clinically, years to decades after initiation and progression of disease. Pathogens and the resulting immune response have been correlated to AD but mechanistic evidence integrating an infection and aging is lacking. Here, we addressed the impact of viral infection on several aspects of AD pathology as a function of age. The data demonstrate that intermittent CMV infection exacerbates age-associated mitochondrial dysfunction and oxidative stress. First, repeated MCMV infection increased basal mitochondrial function over time. Second, mitochondrial specific ROS was elevated in aged, repeated MCMV exposed mice. Third, intermittent infection resulted in significant cognitive impairment. Importantly, the intermittently infected mice, even at the latest timepoint, appear and behave just like the mock infected animals.

We report that intermittent CMV challenge impacts the metabolic function of endothelial cells in the BBB. The degree of functional change increases with viral burden and age. How these changes are mediated continue to be investigated. Metabolic pathways in BMECs are not altered during primary acute infection in young adult mice. Instead, reprogramming occurs incrementally with increased viral burden in advanced age. Inflammatory cytokines have been strongly associated with changes to BBB permeability (Yang et al., 2019). We observed increased inflammatory markers in the blood at 7 DPI, but only minor changes in the brain (Figure 1). In brain lysate, IL-12p40 was significantly increased correlating with published data showing that siRNA knock-down of IL-12p40 in SAMP8 mice ameliorated AD neuropathology (Tan et al., 2014). IL-12(p40) antagonizes IL-12 function and upregulation is needed for efficient MCMV clearance (Carr et al., 1999). CCL5 was also significantly increased and aids in migration of MCMV-specific CD8^+^ T cells (Caldeira-Dantas et al., 2018). CMV encodes a highly specific RANTES decoy receptor (Wang et al., 2004) illustrating the importance of this cytokine during CMV infection. Both cytokines continue to be significantly upregulated 1 MPI, possibly due to paracrine effects of low-level viral persistence or residual viral products. Interestingly, both have been linked to Alzheimer’s disease (Vacinova et al., 2021; Vom Berg et al., 2012). At 1 MPI, we detect low levels of MCMV replication in the salivary gland (Figure 1). IL-1β and TNFα have been implicated in MCMV reactivation *in vivo* (Cook et al., 2006; Simon et al., 2005) possibly establishing a cycle of intermittent reactivation. We did not include assays to quantify MCMV reactivation in this study design, but the significant increase in IgM levels at the 6- and 12-MPI timepoints suggest that MCMV may reactivate more in these cohorts. It has been reported that CMV reactivates more frequently as a function of increased age (Hemmersbach-Miller et al., 2020). Inefficient control of MCMV reactivation by the immune system may explain the emergence of subtle but significant changes at 12 MPI. Surprisingly, cytokine concentrations in the brain are significantly decreased in MCMV infected mice at 12 MPI. Mice were last exposed to MCMV 3 months prior to testing of cytokines so we expected concentrations similar to mock conditions, but not lower concentrations. It is possible that hypoactivation is driving low levels, suggesting dysfunction in one or more glial populations. Alternatively, this may be a response to the delayed expression of inflammatory cytokines seen at 1 MPI. It is also possible that anti-inflammatory mechanisms are over stimulated. Collectively, all scenarios suggest dysregulation of cytokine-associated mechanisms in the brain.

Our -omics approach concurs strongly with the molecular changes we observe. In endothelial cells we measure significant transcriptional changes related to BBB permeability and function. Cadherin-5 (Cdn5), claudin-5 (Cldn5), and CD31 (Pecam1) are crucial for maintaining tight junction integrity and permeability (Ma et al., 2017; Wimmer et al., 2019). Increased expression of CD31 has been linked to neuroinflammation and accumulation of leukocytes (Wimmer et al., 2019). Phospholipid phosphatase 3 (Plpp3) and phospholipase A2 (Pla2g7) are associated with vascular maintenance. Interestingly, phospholipase A2 has recently been shown to regulate phospholipid catabolism during inflammatory and oxidative stress responses (Li et al., 2021; Lv et al., 2021; Spadaro et al., 2022). Decreased expression correlated with lower levels of pro-inflammatory cytokines. The sodium-potassium transporter (ATP1A2) and amino acid transporter (Slc1a3) expression changes support the hypothesis of altered metabolic pathways in brain endothelial cells from MCMV infected mice.

The BBB is the interface between the brain and the periphery and includes BMECs, astrocytes, and pericytes that actively regulate the movement of proteins, nutrients, ions, and small molecules. Systemic inflammation could exert effects on the BBB by affecting the BMECs. Our metabolic profiling of BMECs indicated a significant initial increase in GLUT1 expression in MCMV infected animals which is consistent with previous findings where CMV has been reported to upregulate glycolysis pathways during infection (Combs et al., 2020; Harrison et al., 2022; Munger et al., 2006; Yu et al., 2011). Similar upregulation of glycolysis pathways has been reported in AD brains (Mooradian et al., 1997). However, we noted a significant decrease in GLUT1 expression at 12 MPI. This is an area of active exploration in our lab. GLUT1 is the main mediator of glucose uptake for the brain and alterations have been shown to initiate BBB changes, altering tight junction integrity (Winkler et al., 2015). Reduced GLUT1 may be indicative of reprogrammed BMEC metabolism, shifting dependency to OXPHOS which explains increased basal respiration in 12 MPI mice. This would be similar to a recent report showing CRISPR-Cas9 GLUT1 truncation decreased BMEC bioenergetics and negatively impacted neuro angiogenesis (Pervaiz et al., 2022). Expression of claudin-5, the dominant tight junction protein in the BBB, has been shown to decrease in parallel with GLUT1 following suppression of the Wnt/β-catenin pathway (Wang et al., 2022). Our previous research has demonstrated that CMV infection dysregulates this pathway (Angelova et al., 2012; Zwezdaryk et al., 2016) providing a possible mechanism for these changes.

In addition to the glycolytic changes discussed above, mitochondrial function is significantly altered in MCMV infected BMECs in aged mice, 12 MPI. Our group previously published that CMV infection induces mitochondrial dysfunction in fibroblasts and glioblastoma cell lines (Combs et al., 2020; Harrison et al., 2022). Our current and previous findings are consistent with literature associating metabolism and mitochondrial dysfunction with dementia (Gao et al., 2020; Wilkins and Swerdlow, 2021). VDAC1, the primary channel for the exchange of ADP for ATP through the mitochondrial membrane is intrinsically linked to both the metabolic and the apoptotic functions of mitochondria. The increases in VDAC1 expression and mitochondrial mass with age and infection suggest increased dependence on OXPHOS. Our Seahorse data demonstrate minimal change to mitochondrial ATP production implying OXPHOS is being used for non-bioenergetic purposes. Intriguingly, VDAC1 has been reported to be elevated in the brains of AD patients and in APP transgenic mice (Cuadrado-Tejedor et al., 2011; Manczak and Reddy, 2012). In the context of AD, VDAC1 is required for Aß uptake and apoptosis (Smilansky et al., 2015). Our VDAC1 observations may coincide with hexokinase-influenced glycolytic changes mediating BBB activity (Pastorino et al., 2005). Our model correlates strongly with these studies and demonstrate that further studies exploring VDAC1 function in our model are warranted. Decreased TOMM20 expression was also observed. TOMM20 is responsible for the import of proteins targeted to the mitochondria. A Parkinson’s Disease model provided evidence that overexpression of TOMM20 protected against α-synuclein-induced mitochondrial dysfunction and subsequent neurodegeneration (De Miranda et al., 2020). Based on these observations, decreased TOMM20 levels in our model may suggest impaired Aß clearance due to dysfunctional mitochondrial transport functions.

Age and senescence related decreases in glycolysis and OXPHOS have previously been reported in BMECs suggesting that MCMV infection may accelerate an aging phenotype relative to mock infected animals (Sakamuri et al., 2022). Uncoupling of the ETC may explain the increased proton leak and play a protective role by mitigating ROS production and therefore oxidative damage (Divakaruni and Brand, 2011). Previous research has shown that increased ROS production in the absence of increased ATP production leads to endothelial cell activation and recruitment of immune cells which is supported by our results (Li et al., 2017). Most intriguing is the detection of increased mitochondrial specific superoxide (Figure 4). ROS has been well documented to contribute to neurodegenerative disorders (Valko et al., 2007) specifically neuronal death due to Aß accumulation (Zhang et al., 2009). We measured an increase in total mitochondria suggesting mitochondrial fission is occurring. This is likely due to metabolic stress and can be induced by CMV infection (Combs et al., 2019; Norris and Youle, 2008). It would be interesting to define the impact on mitophagy to understand if damaged mitochondria remaining in the BBB further contribute to BBB functional changes and progression of AD related pathologies.

Astrocytes, the abundant glial cells that reinforce the BMEC tight junctions in the BBB, act as the primary nutrient uptake agents for the brain. Similar to BMECs, astrocytes expressed significantly decreased levels of GLUT1 at 12 MPI. Astrocytic glycolysis is essential for neuron health as glycolysis-derived lactate is taken up by neurons and used in OXPHOS in axon terminals. Studies have demonstrated that astrocytic glycolysis decreases with age and is thought to contribute to AD-related impairments (Yao et al., 2011). Additionally, astrocytic lactate is critical for providing neurons with the energy needed to function properly and form memories (Descalzi et al., 2019). The impact of reduced glucose supply across the BBB into the cerebral parenchyma was not investigated in our model but may play a critical role linking metabolic rewiring to cognitive changes via reduced nutrient availability to neurons. Measuring glucose and lactate concentrations in cortical and hippocampal regions may provide key insights linking intermittent infection with cognitive decline in our model.

The increased presence of T cells in the brain with intermittent infection and age is largely unexplained with our approach but could be driven by increased ROS. We do not detect chronic inflammation in the periphery or the brain. A leaky BBB would provide increased and easier access for T cells to cross the BBB and these changes correlate with infection, but this alone does not fully explain our results. The highly metabolic nature of these T cells (Figure 6) suggest that they may be tissue resident T cells. Murine polyomavirus infection has been reported to produce a set of brain resident memory T cells (Shwetank et al., 2017). Similar results were observed in Vesicular stomatitis virus (VSV) treated mice and it is thought that their persistence in the CNS may be problematic in an environment unaccustomed to immune surveillance (Wakim et al., 2010). However, these viruses are neurotropic and CMV has significant difficulty crossing the BBB in mature immunocompetent mice (Reuter et al., 2004). Future studies will assess BBB permeability in this model.

The most compelling results linking our molecular results with a physical outcome are our cognitive function behavior tests. Foot-shock based passive avoidance test results were very similar between animals at all timepoints indicating the ability to learn fear-based, high salience stimuli effectively. An aging study in BALB/C mice only found significant differences in passive avoidance cognition beginning at 24 months of age (Puglisi-Allegra et al., 1986), so our findings at 12 months were expected. We employed the Y-maze as a measure of spontaneous alternation. Successful alternations between arms indicate an intact short-term recognition memory and by extension suggest intact medial pre-frontal cortex and hippocampal functions. At 1 MPI MCMV-infected mice trend towards a decreased percent of correct alternations in the maze, possible due to lingering sickness behavior (Figure 6) or resulting from the impact of increased pro-inflammatory cytokines in the brain (Figure 1). These events cannot explain the cognitive decline observed at 12 MPI as there is a decreased concentration of pro-inflammatory cytokines (Figure 1). Similarly, age-related activity can be disregarded as both mock and MCMV-infected animals exhibited similar number of alternations in the same timeframe and cognitive performance is determined as a function of total movement in the maze. The significant decrease in cognitive function when merging the cofactors age and infection strongly points to a synergy between CMV infection and age. A study in BALB/c mice was first able to detect cognitive decline using the Y-maze when comparing 4 month old mice to 12 month old mice (Esquivel et al., 2020). The fact that we detected impaired cognition with age matched controls at the 12 MPI timepoint strongly suggests that MCMV infection accelerates cognitive decline.

CMV seropositivity has previously been associated with numerous pathologies including breast and brain cancers (Cobbs et al., 2002; Cui et al., 2018; Harkins et al., 2010), suggesting it may play an oncomodulatory role in disease development and/or progression. Anti-viral therapy targeting CMV has been used to treat patients with glioblastoma and has been shown to increase both overall and progression-free survival (Stragliotto et al., 2020a; Stragliotto et al., 2020b). We have demonstrated that systemic inflammation produces a delayed neuroinflammatory response which may be an initiation event for accelerated cognitive decline. We have linked these changes to altered metabolism in BMECs and enhanced production of ROS suggesting mitochondrial dysfunction and oxidative stress may be driving accelerated disease progression. MCMV is our model pathogen, however, any pathogen, including SARS-CoV-2, which produces transient systemic inflammation, may have the potential to contribute to accelerated cellular aging. It is the additive nature of these pathogen exposures over time which we suspect synergize with myriad other factors to accelerate aging and disease pathology.

Our experimental design includes limitations. We attempted a reductionist approach in which our animals were repeatedly exposed to a single pathogen, very dissimilar to the diverse pathogens experienced throughout life. We also employed a mouse model which is phenotypically “normal” with respect to many of the risk factors strongly correlated with the development of dementia or AD. Our goal was to determine if repeated infection, mimicking the intermittent reactivation or reinfection of CMV that occurs throughout life, without any other complicating factors, could recapitulate some aspects of an AD phenotype. Further, it is unknown, and likely variable, how frequently CMV reactivates in an immunocompetent individual. Our model assumes a single reactivation each decade. For this study, prioritized assessment of structures and processes governing the brain-immune interface so we did not measure Aβ plaques or accumulation of tau tangles, hallmarks of AD. This study also used female mice only, future studies will include males to compare results from current models of dementia and aging (Esquivel et al., 2020). Many of our assays used whole brain or a hemisphere. Despite the generation of exciting data, a region-specific analysis would be of great benefit to further understand how intermittent infection is impacting specific areas of the BBB. Applying this model to transgenic animals predisposed to AD development would provide additional insight into specific cell populations or molecular pathways that change over time. These are all approaches currently being considered by our team.

In summary, our data demonstrate that intermittent CMV infection alters metabolic pathways in the BBB and accelerates cognitive decline. This model makes use of a genotypically and phenotypically normal mouse strain with no predisposition to cognitive deficits or dementia pathology. At 14 months of age, the mice are equivalent to a human in their forties, at which point a human developing dementia would still be in the pro-dromal, symptomless phase. Similar to the course of human disease, MCMV-infected mice are clinically indistinguishable from the controls. Thus our model demonstrates synergy between infection burden and age as potential contributing factors in AD progression. Our work strongly supports a role for pathogens as risk factors in dementia related pathologies. These findings may change the way that we address pathogen exposure and infections especially in populations at a higher risk of developing of dementia.

## AKNOWLEDGEMENTS

The authors would like to thank Constance Porretta of the Flow Cytometry Shared Resource Lab for her assistance in developing and running the multiplexed spectral flow panel. We also would like to thank Robin Kamucheka for her help using De Novo’s FCS Express 7 for our flow analysis and Dr. Raul Freitas, Ms. India Pursell and Ms. Miayla Marcus for technical input. Finally, we would like to thank Dr. Kejing Song from the NextGen Sequencing Core for processing our Multiome samples and the Genomics, Bioinformatics, and Spatial Multiomics Core for help with data analysis.

M.A.A.H was supported by a Louisiana Board of Regents predoctoral fellowship (LEQSF 2017-22-GF-11). P.V.K & R.M. are supported by grants from the NIH, 1R01AG074489 (NIA) and 1R01NS114286 (NINDS). R.M is supported by 1R56AG072676. J.K.K was supported by the Louisiana Board of Regents Endowed Chairs for Eminent Scholars program and PHS grant R35HL139930. E.B.EC is supported by K01MH117343. S.M.J is supported by NIGMS P30GM145498. S.M.J and K.J.Z are supported by NIGMS (P20GM103629). K.J.Z and E.B.EC are supported by pilot grants from the Infectious Disease Society of America (IDSA) and the Tulane Brain Institute.

## AUTHOR CONTRIBUTIONS

Conceptualization, M.A.A.H., E.B.EC., and K.J.Z.; Methodology, M.A.A.H. and K.J.Z.; Formal Analysis, M.A.A.H., M.J.J., M.M.H. M.S.K., and S.K., K.J.Z.; Investigation, M.A.A.H., S.L.M., G.A.R., D.J.R., C.H.M., S.SVP.S., H.W., L.P.G, M.J.J., and E.B.EC.; Resources, K.J.Z., E.B.N., C.S., and J.K.K.; Data Curation, M.A.A.H. and K.J.Z.; Writing, M.A.A.H. and K.J.Z.; Review and Editing, S.L.M., S.SVP.S., S.K., J.K.K., S.M.J., R.M., and E.B.EC; Visualization, M.A.A.H., M.M.H., M.S.K., S.K., E.B.EC, K.J.Z; Supervision, K.J.Z., S.K., R.M., and P.K.; Project Administration, K.J.Z.; Funding Acquisition K.J.Z. and S.M.J.

## VIRUS PROPAGATION AND QUANTIFICATION

Murine cytomegalovirus (MCMV) Smith strain was passaged through mice and quantified via plaque assay as previously described (Brune et al., 2001; Zurbach et al., 2014). Briefly, mice were

i.p. injected with MCMV and sacrificed at 14 days post infection. Salivary glands were homogenized in PBS, aliquoted, and stored at -80°C. Virus was quantified by plaque assay using M2-10B4 cells and a viscous overlay followed by fixation with formaldehyde and staining with 0.02% methylene blue.

## ANIMALS, INFECTIONS, AND SAMPLE COLLECTIONS

Female BALB/cJ mice aged 6 weeks were obtained from Charles River and housed in disposable, sterile cages in groups of 5 with food and water provided *ad libitum*. All animal experiments were approved by the Tulane University Institutional Animal Care and Use Committee. MCMV infections were conducted via i.p. injection with 5x10^4^ plaque forming units in 100uL sterile PBS. They were maintained on a diurnal light cycle (8-6). Mice were euthanized via i.p. injection with a sub-lethal dose of ketamine and xylazine followed by exsanguination. Blood collected via cardiac puncture was processed for plasma. Mice were then perfused with ice-cold PBS and tissues were collected and processed as needed.

## BEHAVIORAL TESTING

Behavior in mice was evaluated using the Y-maze, Passive Avoidance, and Hot Plate tasks at 1 month post initial infection or 12 months post initial infection. The behavior tests described below were performed in the order they are listed. All behavioral testing was performed between 0800 and 1400 hrs. All animals underwent testing by a pair of treatment blinded researchers. Peroxiguard was used to disinfect and deodorize the apparatuses between each mouse trial.

### Y-Maze

The Y-Maze was employed as a measurement of spontaneous alternation/short-term recognition memory (Swonger and Rech, 1972). Mice were placed in the three-armed maze with arm length of 38 cm, width of 8.25 cm, and height of 13.25 cm facing the end of arm designated “A”. Mice were allowed to freely explore the maze for 8 minutes. Movement around the maze was recorded by an overhead camera and tracked using ANY-Maze 7.1 software. Mice were provided with large black geometric spatial clues on white backgrounds to help them navigate and discriminate between arms. Spontaneous alternation was calculated as percentage of correct alternations between the three arms without revisiting the previous arm. Total alternation (number of total arm entries minus 2).

### Passive Avoidance

As a measure of single-trial fear based aversive learning, mice were subjected to the step-through inhibitory (passive) avoidance task (Jarvik and Kopp, 1967). In this assay mice are placed in a two-chamber device with a guillotine door between them. One chamber is brightly lit while the other is dark. The floor of both chambers contains metal rungs on the floor connected to a shock delivery apparatus (Maze Engineers). For the training portion of the test, mice were placed in the brightly illuminated side (approx.. 1300 lux) and allowed to explore until they entered the dark chamber. Once the mouse entered the dark chamber the door was closed and the mouse was administered a 2 second foot shock (0.25 mA). Mice that did not enter the dark chamber within 120 seconds were gently pushed to the dark chamber and a foot shock was administered. After the training trial was completed, animals were tested for learning ability (10 minutes after training trial) and memory retention (24 hours later) by placing the mice in the bright chamber and recording latency to enter dark chamber. These retention trials were limited to 300 seconds.

### Hot Plate

To examine nociceptive ability, mice were exposed to the hot plate test (Woolfe and Macdonald, 1944). The hot plate (Maze Engineers) was set to 55.0°C and the mouse was placed onto the plate surface. Latency to first nociceptive behavior as well as total number of nociceptive behaviors was recorded for the 30 second duration of the experiment. Nociceptive behaviors were defined as shaking or flicking of hindlimbs and jumping.

## SINGLE CELL MULTIOME ATAC AND GENE EXPRESSION SEQUENCING

### Nuclei Isolation

Nuclei isolation for multiome sequencing was performed following protocol revision D (10X Genomics) with minor modifications. Following euthanasia and perfusion, the brain was extracted and flash frozen in liquid nitrogen. The brain was lysed with 0.1x lysis buffer and homogenized with scissors and a pellet pestle. Homogenate was then incubated for 3 minutes on ice and lysis stopped via the addition of wash buffer. The suspension was filtered through a 70 µm filter and then subjected to a 30% Percoll Plus centrifugation to remove myelin. Nuclei were washed and resuspended in nuclei buffer. Finally, nuclei were visualized on a cell counter for nuclear membrane integrity (lack of blebbing) and counting.

### Library Prep and Sequencing

The Chromium Next GEM Single Cell Multiome ATAC and GEX protocol (CG000338) and 10x reagent kit were used to prepare ∼10,000 nuclei/sample for the Multiome sequencing. Briefly, nuclei suspensions were incubated with a Transposon Mix and DNA fragments were transposed and barcoded. The transposed nuclei were partitioned into nanoliter-scale Gel Bead-In EMulsion (GEMs). The barcoded transposed DNA and barcoded full-length cDNA from poly-adenylated mRNA were generated and amplified by PCR. P7 and a sample index were added to transposed DNA during ATAC library construction via PCR. Barcoded cDNA enzymatic fragmentation, end-repair, A-tailing, and adaptor ligation were followed by PCR amplification. Both final ATAC libraries and 3′ Gene Expression libraries generated contained standard Illumina P5 and P7 paired-end constructs. Library quality controls were performed by using an Agilent High Sensitivity DNA kit with Agilent 2100 Bioanalyzer and quantified by Qubit 2.0 fluorometer. Libraries were sequenced separately with individual parameter settings. Pooled libraries at 650 pM were sequenced with paired end dual index configuration on an Illumina NextSeq 2000 using Illumina P3 100 cycle kit. Cell Ranger arc version 2.0.0 (10X Genomics) was used to process raw sequencing data and mapped to mouse genome mm10-2020-A to generate differentially expressed genes between specified cell clusters represented by Loupe Cell Browser (10X Genomics).

### Multiome Data Analysis

Cell Ranger ARC v2.0 (10x Genomics) was used to process raw sequencing data for further downstream analyses. After filtering, we used 27,798 cells with high quality transcriptome and epigenome profiles. We followed the algorithms Seurat (v4.0) R package (v4.0.2) to generate the uniform manifold approximation and projection (UMAP) embeddings and differentially expressed (DE) gene. Fragment files were used as input for Seurat analysis. The input fragment file was processed as chunks per chromosome and stored in HDF5 format (hierarchical data format version 5), allowing rapid access and efficient read and write functions during analysis. The quality control steps involved the removal of all low-quality cells and predicted heterotypic doublets from our analysis. We used Morpheus (https://software.broadinstitute.org/morpheus/) software to generate the top 30 DE genes heatmap. A relative color scheme uses the minimum 0 and maximum 0.1 values in each row to convert values to colors.

## RNA ISOLATION AND qRT-PCR

Salivary glands were flash frozen in liquid nitrogen and stored at -80°C. RNA was isolated from 30 mg of salivary gland using the RNeasy minikit from Qiagen. iScript cDNA synthesis kit was used to created cDNA from 1 ug of RNA. qRT-PCR was conducted on 50 ng of each sample using Sso Advanced SYBR Green Supermix and primers for MCMV immediate early gene 1 (UL123) and murine 36B4 which was used as a housekeeping gene. qPCR products were run on a 2% agarose gel to confirm product size.

## FLOW CYTOMETRY

Brains were processed for flow cytometry as previously described (Calvo et al., 2020). Briefly, they were incubated with 5mg collagenase II and homogenized in MACS C-tubes with 5 mg collagenase II and 100U DNAse I in HBSS with 3mM calcium chloride. Myelin debris was removed via a 30% Percoll Plus gradient. Cells were washed and resuspended in PBS with 0.4% BSA.

### Neuro Flow Panel

Cells were incubated with fixable live dead stain and Fc block for 15 minutes at RT. Following washes with 20% BD Horizon Brilliant Stain Buffer, cells were incubated in an extracellular antibody cocktail for 20 minutes at RT. Cells were washed, fixed and permeabilized with eBioscience FoxP3 Transcription Factor staining kit for 45 minutes. Fixed cells were washed and incubated with intracellular antibodies for 45 minutes at room temperature. Following washes, cells were resuspended in PBS with 0.5% BSA and flow cytometry was conducted on a Cytek Aurora. Data was analyzed using FCS Express 7.14.

### Live Cell Mitochondrial Panel

Cells were incubated in a cocktail of anti-CD31 BV605 antibody at 1:100, 1 μM nonyl acridine orange (mitochondrial mass), 2.5 μM MitoSOX Red (superoxide production), and 250 nM MitoTracker Red FM (mitochondrial membrane potential) for 45 minutes at 37°C. Following washes in PBS, cells were resuspended in PBS with Sytox Blue dead cell stain, incubated for 15 minutes, and flow cytometry was performed on a BD LSRFortessa. Data was analyzed using FCS Express 7.14.

## CYTOKINE ANALYSIS

Mouse brains were minced briefly in BioRad Cell Lysis Buffer followed by homogenization in MACS S-tubes and freezing at -80°C. Following thaw, lysate was sonicated briefly and then centrifuged to pellet debris before being quantified by BCA. Brain lysate was diluted to 4mg/mL in Bio-Plex diluent.

Both plasma and brain lysate cytokine levels were assessed using a Bio-Plex murine cytokine 23-plex assay kit at a dilution of 1:2 according to manufacturer recommendations. Briefly, samples were added to a 96-well plate with magnetic beads and incubated for 30 minutes while shaking. Well, were washed three times and followed by addition of detection antibodies for 30 minutes at room temperature. Three more washes were conducted and then streptavidin-PE was added and incubated for 10 minutes. After another three washes samples were read on the Bio-Plex 200 system with HTF. Standard curves were created for each cytokine from 1-32,000 pg/mL, a five-parameter logistic regression was fit, and sample cytokine concentrations were interpolated.

## ELISA

Antigen-specific quantitative ELISAs were performed using similar methods as described in ref. (Norton et al., 2015). Briefly, 96-well plates were coated with 0.25 μg/well of UV-inactivated MCMV. Serial dilutions of plasma from animals were incubated for an hour at room temperature followed by detection using AP-conjugated rabbit anti-mouse IgG or IgM. ELISAs were quantified using dilutions of purified mouse standards IgG1-κ or IgM-κ and the results were expressed as ELISA units/mL (EU/mL).

## BRAIN MICROVESSEL ISOLATION

Excised brains were stored in ice-cold PBS. Brain microvessels (BMVs) were isolated as previously described {Sakamuri, 2022 #1160;Sakamuri, 2022 #1218}. Briefly, hemispheres were rolled on filter paper to remove surface vessels. Brains were homogenized in PBS, centrifuged, and the pellet was resuspended in 17.5% dextran solution. This process was repeated, and each time floating white matter was removed from the pellet prior to resuspension. The resuspended vessels were serially filtered through a 300 μm and 40 μm filters and then the BMV pellet is resuspended in Seahorse XF DMEM media.

## SEAHORSE

BMVs were loaded into Seahorse XFe24 cell plates with one mouse used for each well. Plates were centrifuged and incubated for 1 hour at 37°C in a non-CO_2_ incubator. Mitochondrial function was assessed using the Mito Stress Test Kit. Basal media contained 25 mM glucose, 10 mM sodium pyruvate, and 2 mM glutamate. The assay was completed using the manufacturers recommended protocols with following injections: Oligomycin (5 μM), FCCP (5 μM), and Rotenone/Antimycin A (2 μM/10 μM). Total protein per well was determined via BCA and oxygen consumption rates were normalized to total protein for each well. Normalized OCR was used to determine basal and maximal respiration, ATP coupled respiration, proton leak, and spare respiratory capacity.

## STATISTICAL ANALYSIS

Statistical analyses were performed using Graphpad Prism v9. All data are expressed as the mean plus standard error of the mean (SEM) from at least three independent experiments or animals. Comparisons between groups were assessed using unpaired *t*-tests, ordinary one-way ANOVA, or repeated measures ANOVA followed by Dunett’s multiple comparisons test where appropriate. Outliers were identified and excluded where necessary using the ROUT method with a maximum desired false discovery rate (Q) of 2%. For all comparisons, a *p* < 0.05 was considered significant.

## KEY RESOURCES TABLE

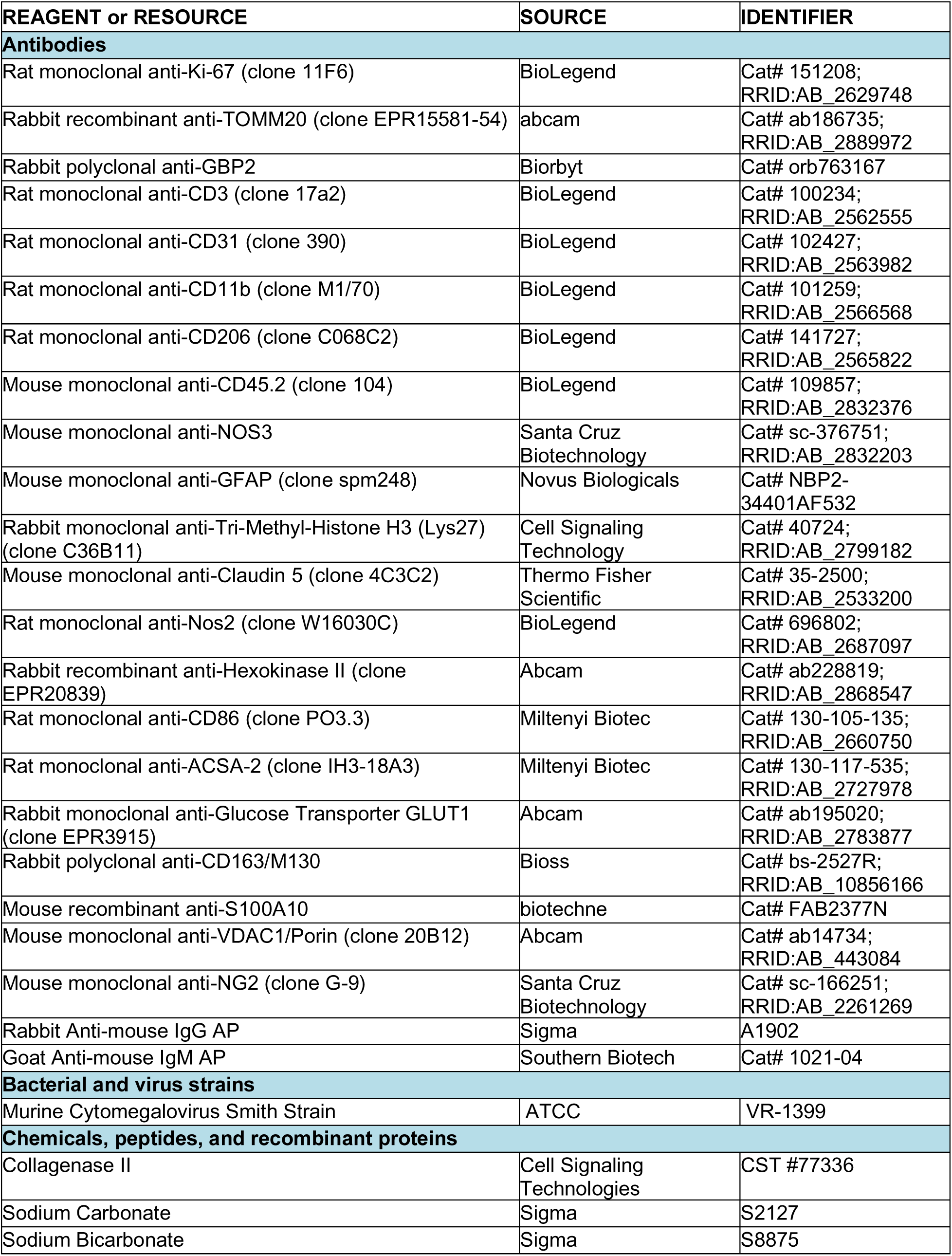
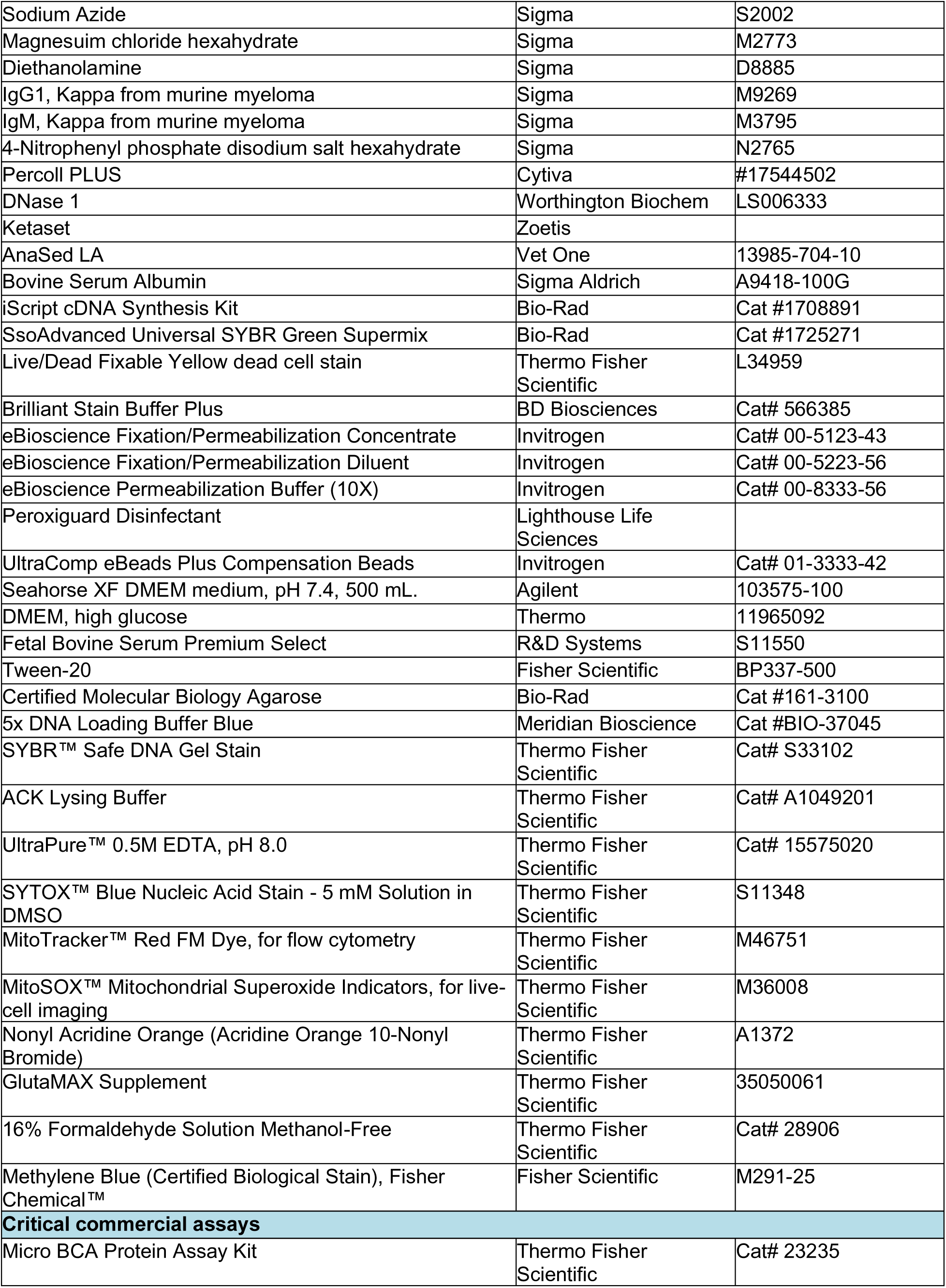
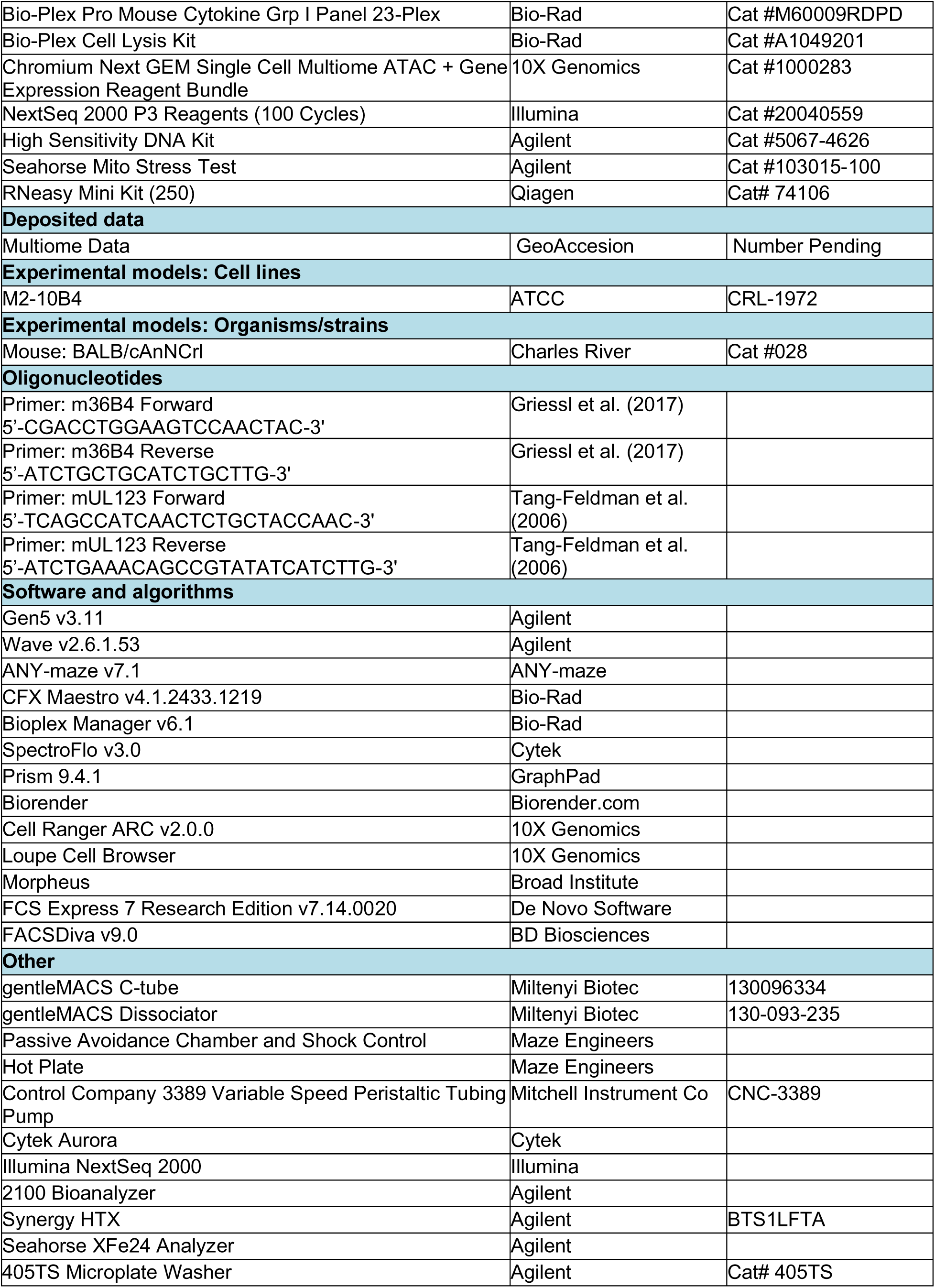
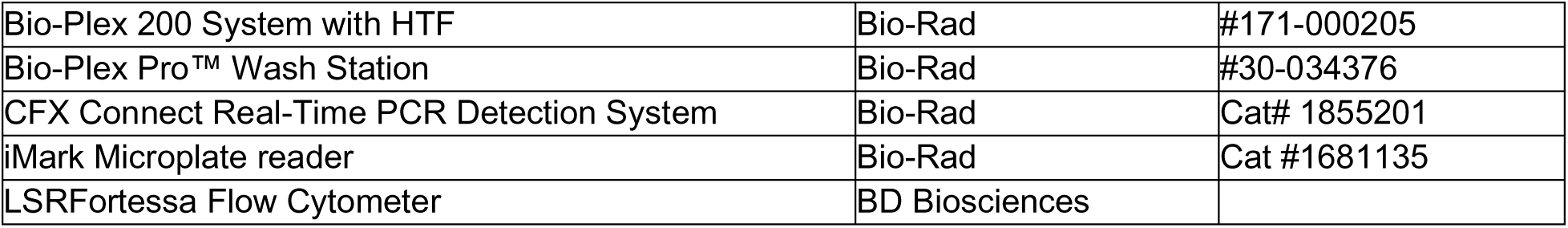

---

**Supplemental Figure 1:**
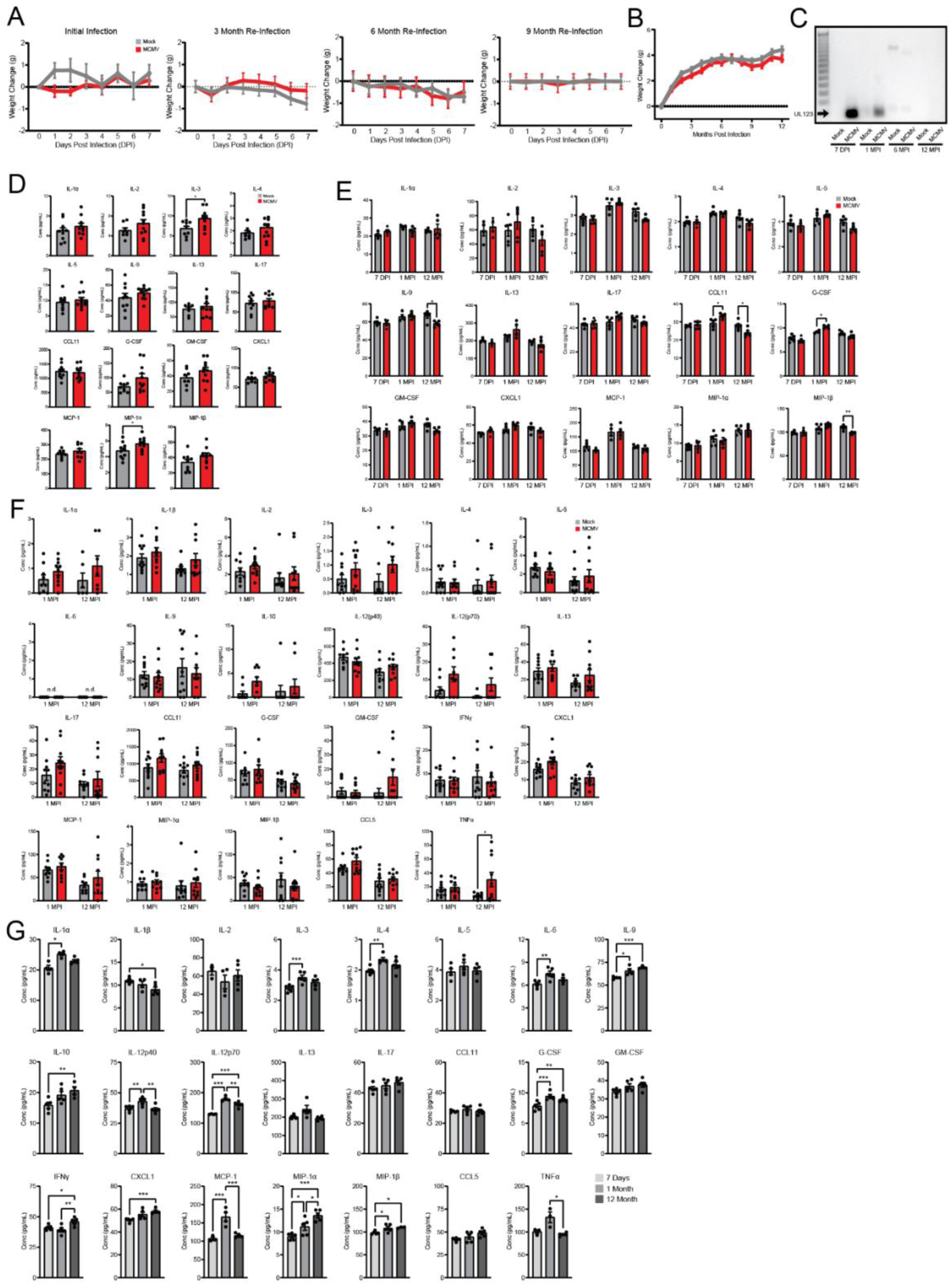
Intermittent MCMV infection exhibits delayed inflammatory cytokine expression in the brain. (A) Weight change in mock- and MCMV-infected mice over 7 DPI from initial infection through the 9-month infection time point. (B) Monthly weight of animals from initial infection through 12 MPI. (C) Agarose gel displaying UL123 qRT-PCR products from salivary glands at 7 DPI, 1 MPI, 6 MPI, and 12 MPI. Cytokine levels from (D) plasma at 7 DPI and (E) brain homogenate at 7 DPI, 1 MPI, and 12 MPI as measured by a multiplex immunoassay. Cytokine levels in (F) plasma at 1 and 12 MPI, and (G) brain homogenate in mock infected animals at 7 DPI, 1 MPI, and 12 MPI. Graphs represent pooled data from at least three individual animals. Mean ± the SEM. *, *p* < 0.05; **, *p* < 0.01; ***, *p* < 0.001. DPI = days post infection, MPI = months post infection.

**Figure S2:**
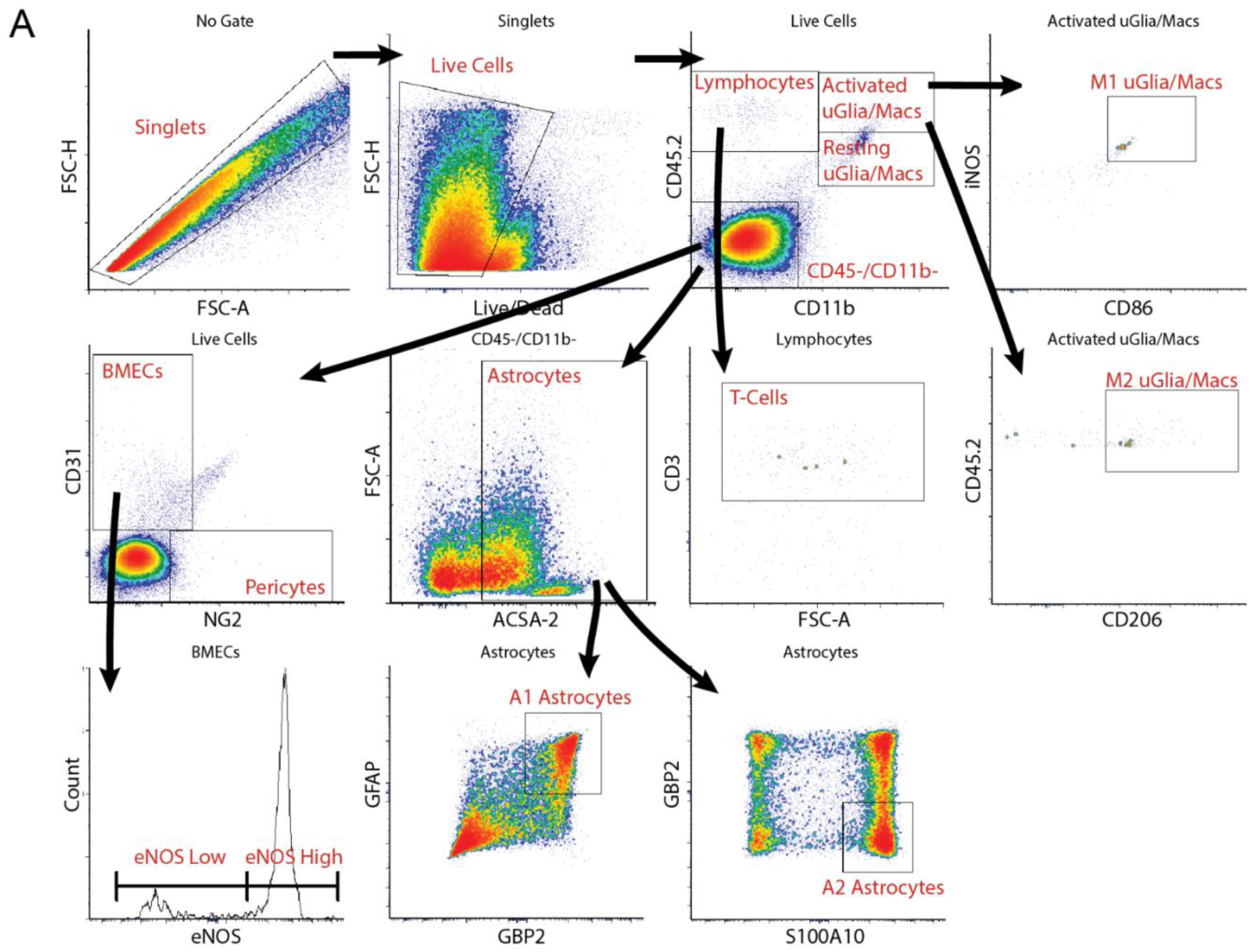
Brain phenotyping flow cytometry panel gating strategy.

**Figure S3:**
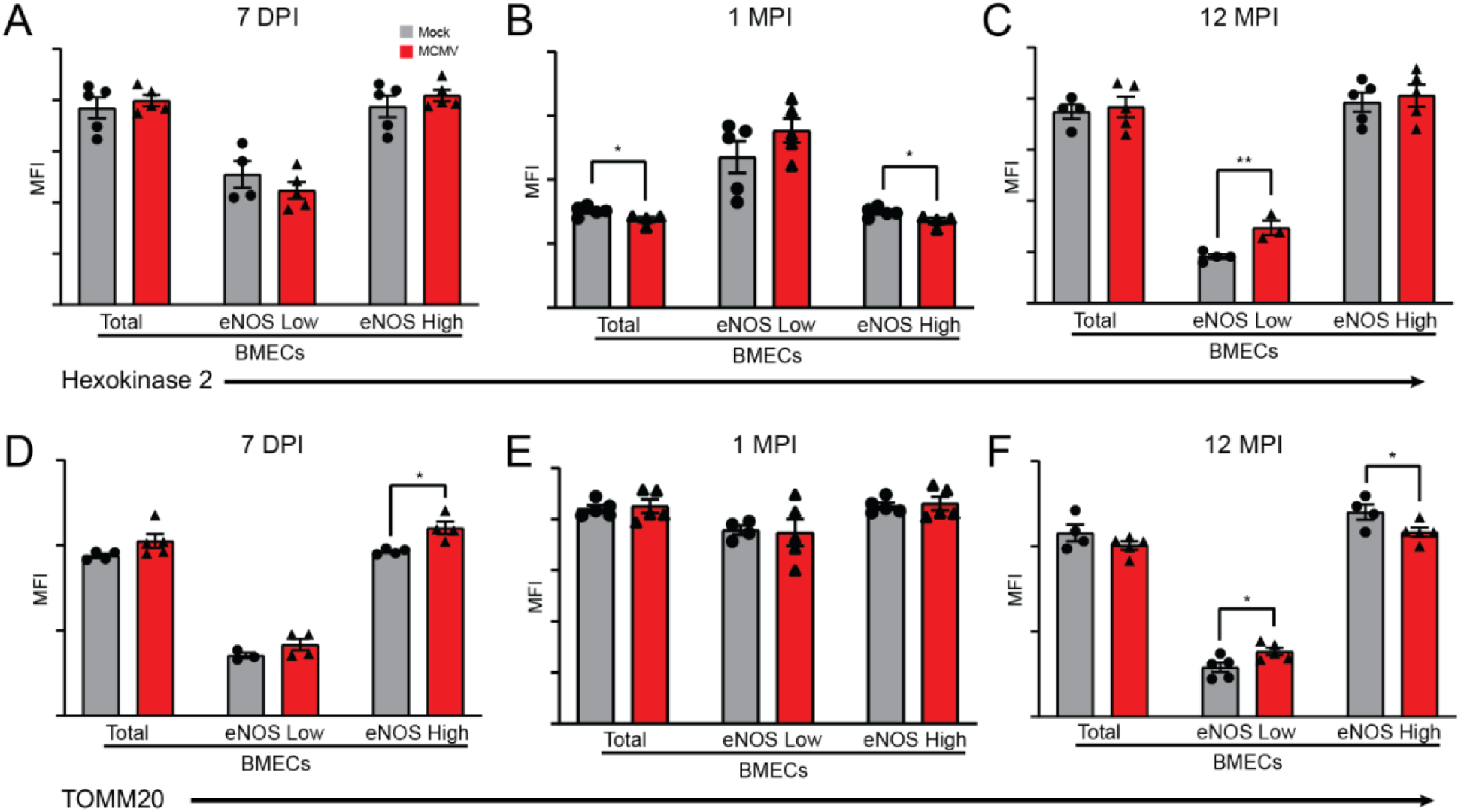
Blood brain barrier metabolic pathways are rewired with increased MCMV exposure. Flow cytometry was used to examine expression of Hexokinase 2 in populations of BMECs at (A) 7 DPI, (B) 1 MPI, and (C) 12 MPI. Expression of TOMM20 in populations of BMECs at (D) 7 DPI, (E) 1 MPI, and (F) 12 MPI. Graphs represent pooled data from at least three individual animals. Mean ± the SEM. *, *p* < 0.05; **, *p* < 0.01; ***, *p* < 0.001. eNOS = endothelial nitric oxide synthase, MFI = mean fluorescent intensity.

**Figure S4:**
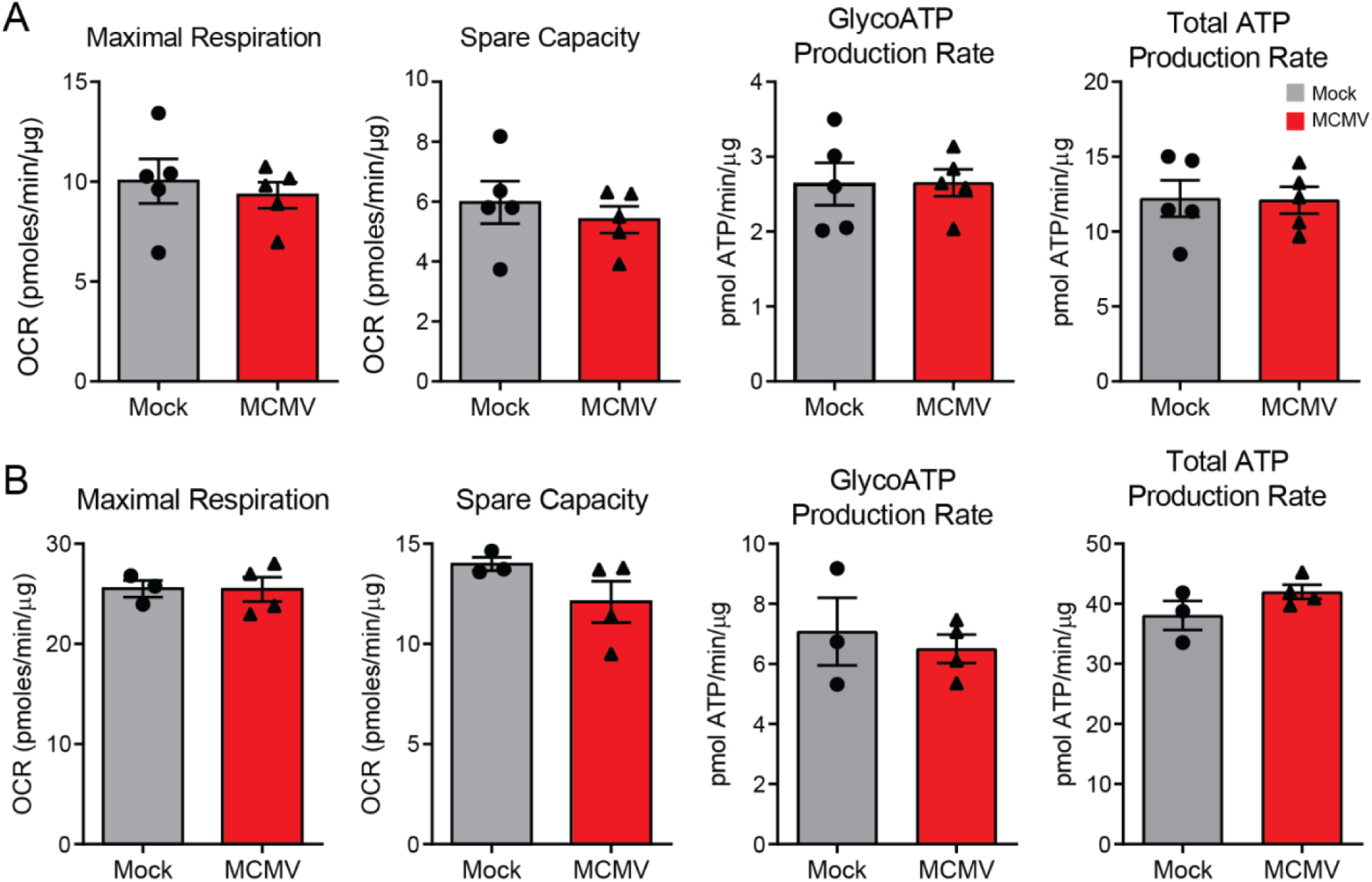
Hallmarks of mitochondrial dysfunction and oxidative stress are elevated in BMECs intermittently exposed to MCMV. Maximal Respiration, Spare Respiratory Capacity, GlycoATP Production Rate, and Total ATP Production Rate were derived from the (S4A) 1 MPI and (S4B) 12 MPI measurements. Graphs represent pooled data from at least three individual animals. Mean ± the SEM. OCR = oxygen consumption rate.

**Figure S5:**
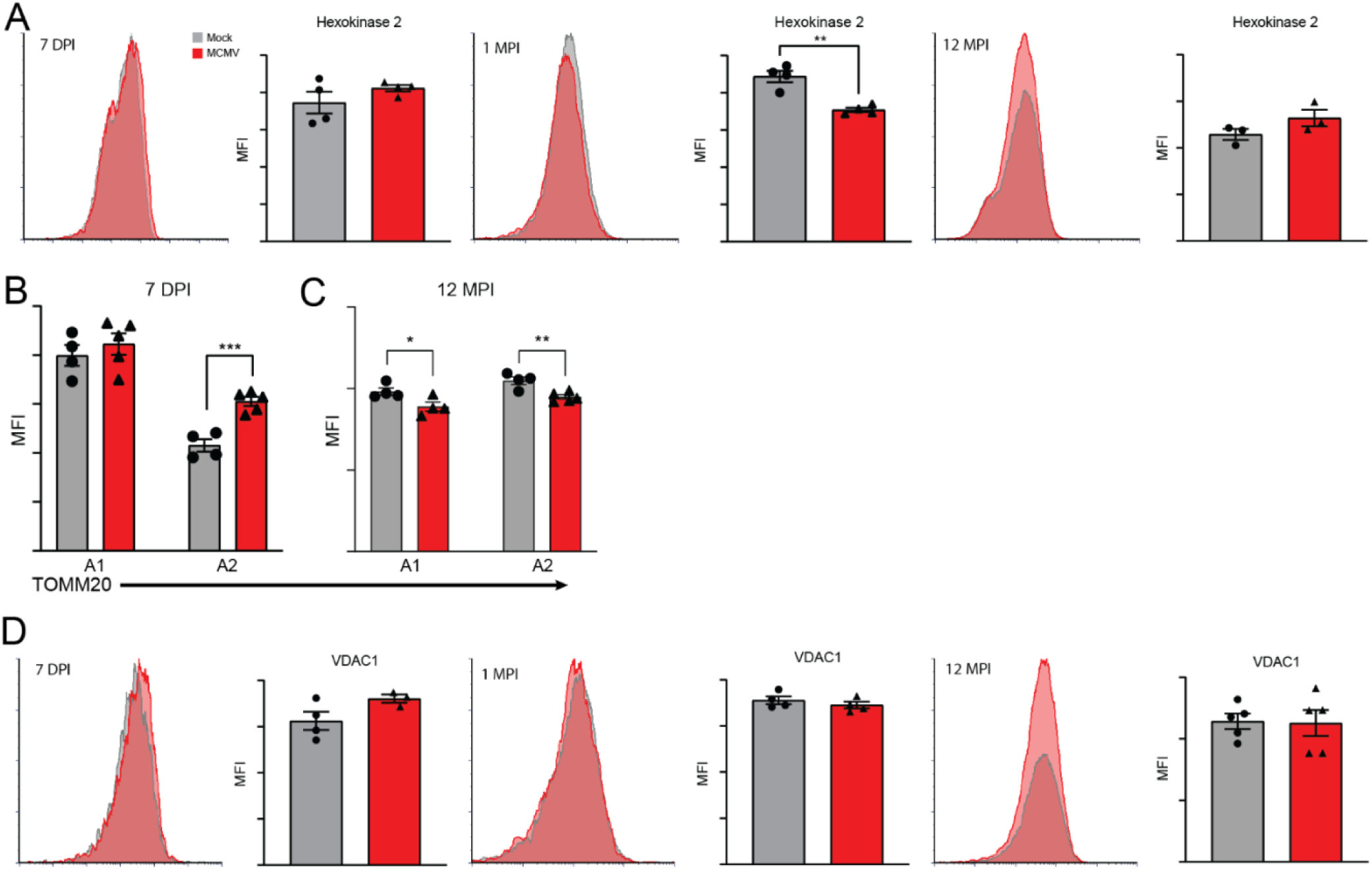
MCMV infection metabolically rewires astrocytes as a function of age. Flow cytometry expression of (A) Hexokinase 2 in astrocytes at 7 DPI, 1 MPI and 12 MPI. TOMM20 expression in A1 and A2 astrocytes at (B) 7 DPI and (C) 12 MPI. (D) VDAC1 expression in astrocytes at 7 DPI, 1 MPI, and 12 MPI. Graphs represent pooled data from at least three individual animals. Mean ± the SEM. *, *p* < 0.05; **, *p* < 0.01; ***, *p* < 0.001. DPI = days post infection, MPI = months post infection, MFI = mean fluorescent intensity.

**Figure S6:**
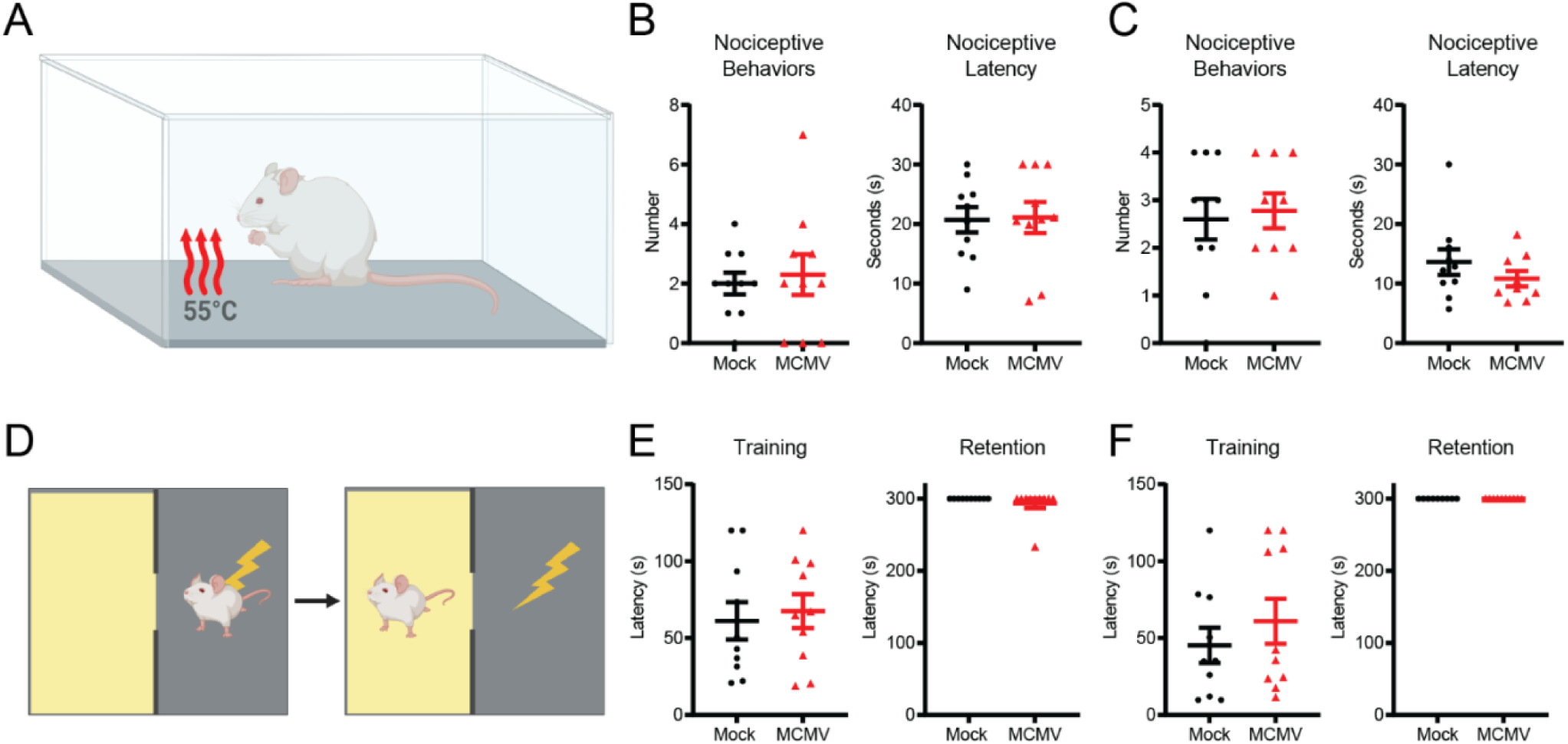
Cognition is impaired with intermittent MCMV infection. (A) Cartoon of the hot plate assay. Quantification of the number and latency to produce nociceptive behaviors in the hot plate assay at (B) 1 MPI and (C) 12 MPI. (D) Cartoon of the passive avoidance foot shock assay. Quantification of latency to enter the dark, foot-shock chamber during training and 24-hour retention testing in (E) 1 MPI (F) 12 MPI animals. Graphs represent pooled data from at least three individual animals. Mean ± the SEM. *, *p* < 0.05; **, *p* < 0.01; ***, *p* < 0.001.

**Table S1:**
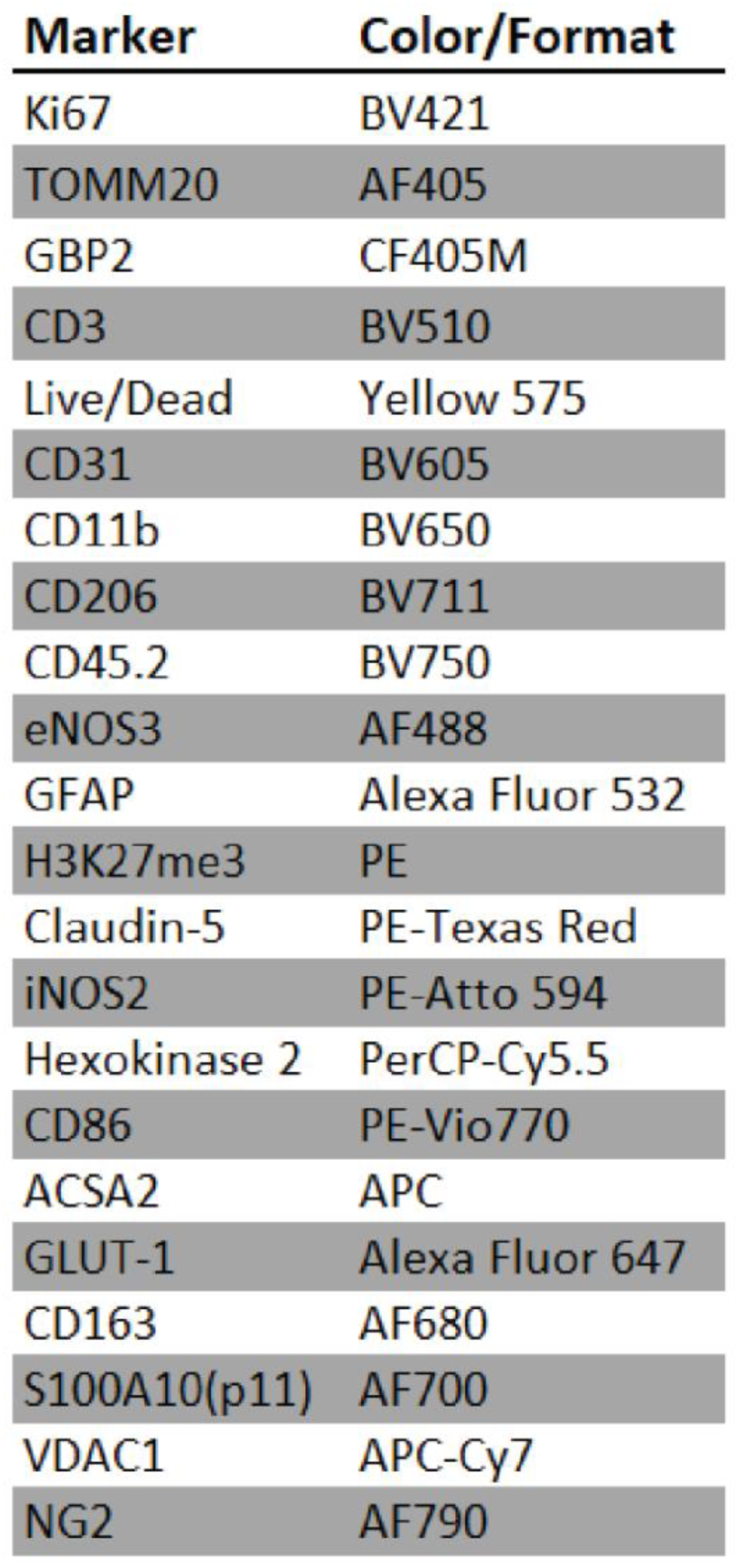
Antibodies and fluorophores.

| **Marker** | **Color/Format** |
| --- | --- |
| Ki67 | BV421 |
| TOMM20 | AF405 |
| GBP2 | CF405M |
| CD3 | BV510 |
| Live/Dead | Yellow 575 |
| CD31 | BV605 |
| CD11b | BV650 |
| CD206 | BV711 |
| CD45.2 | BV750 |
| eNOS3 | AF488 |
| GFAP | Alexa Fluor 532 |
| H3K27me3 | PE |
| Claudin-5 | PE-Texas Red |
| iNOS2 | PE-Atto 594 |
| Hexokinase 2 | PerCP-Cy5.5 |
| CD86 | PE-Vio770 |
| ACSA2 | APC |
| GLUT-1 | Alexa Fluor 647 |
| CD163 | AF680 |
| S100A10(p11) | AF700 |
| VDAC1 | APC-Cy7 |
| NG2 | AF790 |

